# Multi-species genome-wide CRISPR screens identify conserved suppressors of cold-induced cell death

**DOI:** 10.1101/2024.07.25.605098

**Authors:** Breanna Lam, Kathrin M. Kajderowicz, Heather R. Keys, Julian M. Roessler, Tara G. Thakurta, Evgeni M. Frenkel, Adina Kirkland, Punam Bisht, Mohamed A. El-Brolosy, Rudolf Jaenisch, George W. Bell, Jonathan S. Weissman, Eric C. Griffith, Sinisa Hrvatin

## Abstract

Cells must adapt to environmental changes to maintain homeostasis. One of the most striking environmental adaptations is entry into hibernation during which core body temperature can decrease from 37°C to as low as 4°C. How mammalian cells, which evolved to optimally function within a narrow range of temperatures, adapt to this profound decrease in temperature remains poorly understood. In this study, we conducted the first genome-scale CRISPR-Cas9 screen in cells derived from Syrian hamster, a facultative hibernator, as well as human cells to investigate the genetic basis of cold tolerance in a hibernator and a non-hibernator in an unbiased manner. Both screens independently revealed glutathione peroxidase 4 (GPX4), a selenium-containing enzyme, and associated proteins as critical for cold tolerance. We utilized genetic and pharmacological approaches to demonstrate that GPX4 is active in the cold and its catalytic activity is required for cold tolerance. Furthermore, we show that the role of GPX4 as a suppressor of cold-induced cell death extends across hibernating species, including 13-lined ground squirrels and greater horseshoe bats, highlighting the possibility of evolutionary conservation of this mechanism of cold tolerance. This study identifies GPX4 as a key modulator of mammalian cold tolerance and advances our understanding of the evolved mechanisms by which cells mitigate cold-associated damage—one of the most common challenges faced by cells and organisms in nature.

## Introduction

Rapid temperature changes pose a challenge to all clades of life, including endotherms. Homeotherms such as mammals routinely defend a core body temperature set-point, with profound thermal deviations leading to organ dysfunction and death^1–4^. Indeed, several studies have shown that while homeotherm-derived cells typically possess a capacity for mild cold tolerance in culture, primary cultured cells and cell lines derived from a number of mammalian species, including humans, exhibit high rates of cell death following prolonged severe cold exposure^5–8^. Although considerable work has focused on understanding the cellular responses and adaptations to an increased temperature (heat stress and heat-shock)^9–11^, little is known about how cell biological processes modulate cell sensitivity to decreased temperatures.

The mechanisms underlying cold-induced death remain poorly understood but have been hypothesized to involve phase transition of cellular membranes, with the resulting membrane damage leading to ion gradient disruption and irrecoverable cellular dysfunction^12–15^. However, many warm-blooded animals, including some primates, have evolved the ability to enter torpor and hibernation – states during which body temperature can decrease far below its homeostatic set-point^16–18^. In many animals, *torpor* is a state of profoundly reduced metabolic rate and body temperature lasting from hours to days, while *hibernation* is a seasonal behavior comprising multiple bouts of *torpor* interrupted by periodic arousals to normal body temperature. During torpor, the core body temperature of many small hibernators including the Syrian golden hamster (*Mesocricetus auratus*), the 13-lined ground squirrel (*Ictidomys tridecemlineatus*), and the greater horseshoe bat (*Rhinolophus ferrumequinum*), reaches 4-10°C^19–21^. Animals can remain at these low temperatures for extended periods of time (up to several weeks), indicating that their cells and tissues are either genetically predisposed to cold tolerance, or that they can adapt to tolerate long-term cold exposure^19–22^.

Although in principle the ability to tolerate cold temperatures could be conveyed by systemic factors present in torpid or hibernating animals, cell lines derived from hibernating rodents maintain high viability when exposed to 4°C for several days, indicating the presence of cell-intrinsic mechanisms of cold tolerance^5–8^. Several studies have reported distinctive responses of hibernator-derived cells in the cold. For example, Syrian hamster-derived epithelial-like hamster kidney (HaK) cells maintain mitochondrial membrane potential and ATP production during cold exposure^7^, and both HaK and epithelial-like smooth muscle cells derived from the Syrian golden hamster are able to prevent excess reactive oxygen species damage in response to cold exposure^5^. Similarly, unlike human induced pluripotent stem cell (iPSC)-derived neurons, cultures of iPSC-derived neurons from 13-lined ground squirrels retain microtubule stability in the cold^6^. Despite metabolic and cell biological differences between various hibernator and non-hibernator-derived cells in the context of cold exposure, and several identified modulators of acute cold stress such as forkhead box O1 (FOXO1) and 5-aminolevulinate (5-ALA)^5,6,23,24^, the underlying genetic modulators of cold tolerance have yet to be systematically explored. Additionally, it is still unknown to what extent the pathways that regulate cold tolerance in one hibernating species are conserved across other evolutionarily distant hibernators and potentially even present in non-hibernators.

Here, we carry out the first unbiased genome-scale CRISPR-Cas9 screens in cells derived from a cold-tolerant hibernator (Syrian hamster) as well as human (non-hibernator) cells to identify genetic pathways required for long-term cold tolerance. Surprisingly, these screens independently identify a common mechanism dependent on the selenocysteine-containing enzyme Glutathione Peroxidase 4 (GPX4) as a key mediator of cellular cold tolerance. Employing genetic and pharmacological approaches, we confirm these findings and demonstrate that increased GPX4 activity is sufficient to improve cold tolerance in human cells. We further show that functional GPX4 is required for cold tolerance across several hibernating and non-hibernating mammals, including in distantly related hibernating horseshoe bats (*Rhinolophus ferrumequinum*), and propose that the GPX4 pathway may be a widespread, evolutionarily ancient metazoan mechanism of cold tolerance across endotherms.

## Results

### Assay for measuring viability of cold-exposed cells

Our initial evaluation of the ability of human K562 leukemia cells to tolerate prolonged exposure to extreme cold (4°C) temperatures led us to establish a new method for assessing viability of cold-exposed cells. To avoid confounding changes in media composition or pH, we transferred cultured cells into incubators cooled to 4°C with the CO_2_ concentration calibrated to maintain physiological pH (7.4). Cell survival was assessed based on exclusion of trypan blue, a membrane-impermeable dye, immediately upon removal from 4°C. We observed a 27 ± 3% survival rate for K562 cells following 24 hours at 4°C using this approach. However, recounting live cells after a 24-hour rewarming at 37°C revealed a ∼6-fold increase in live cells, inconsistent with the reported 16-24 hour doubling rate of this cell line (**Figure 1b, Figure S1a**). Notably the rate of cell division was indistinguishable between cells that were continuously maintained at 37°C and cells rewarmed to 37°C following 24 hours at 4°C (**Figure S1b**), further confirming the viability of rewarmed cells. We reasoned therefore that a substantial fraction of trypan blue-positive cells at 4°C remain alive and capable of cell division and recovery upon rewarming, and carried out a time course of trypan blue staining shortly after removal from 4°C (**Figure 1a**). We found that the fraction of trypan blue-positive cells dramatically reduced following a brief rewarming period (30 minutes to 1 hour, either at room temperature or at 37°C), consistent with the number of live cells observed after 24 hours of rewarming (**Figure 1b, c, Figure S1a,c**). To determine whether this effect was isolated to K562 cells, we measured trypan blue-positive cells at 4°C or following a brief period of rewarming in several human cell lines (HEK 293T, HeLa, and RPE1 cells), as well as Syrian hamster kidney fibroblasts (BHK-21 cells) (**Figure S1d-g**). We consistently observed that, absent a rewarming period, trypan blue staining significantly overestimates cold-induced cell death. Moreover, we observed the same phenomenon of transient, reversible cold-induced membrane permeability using an orthogonal flow cytometry-based assay we developed, which utilized two DNA-binding dyes, TO-PRO-3 and DAPI. Both dyes are normally membrane-impermeable and can only enter cells whose plasma membranes are compromised. In this assay, TO-PRO-3 was added at 4°C to label any cell that was permeable during cold exposure. Cells were then rewarmed to 37°C for 1 hour, after which DAPI was added to identify cells that are dye impermeable at 37°C, which we interpreted as live cells. We tested this orthogonal assay in K562, RPE1, and BHK-21 cells incubated at 4°C for 24 hours and found in all cell lines that TO-PRO-3 fluorescent intensity was significantly higher in live cells exposed at 4°C relative to 37°C, confirming that the cold-induced dye permeability in live cells (**Figure S2)**. Given that our flow cytometry and trypan blue counting assays converged on similar results, we thus adopted a brief rewarming period (∼30-minutes) prior to trypan blue incubation as a standard assay to measure cell viability of cold-exposed cells in our subsequent studies.

**Figure 1.**
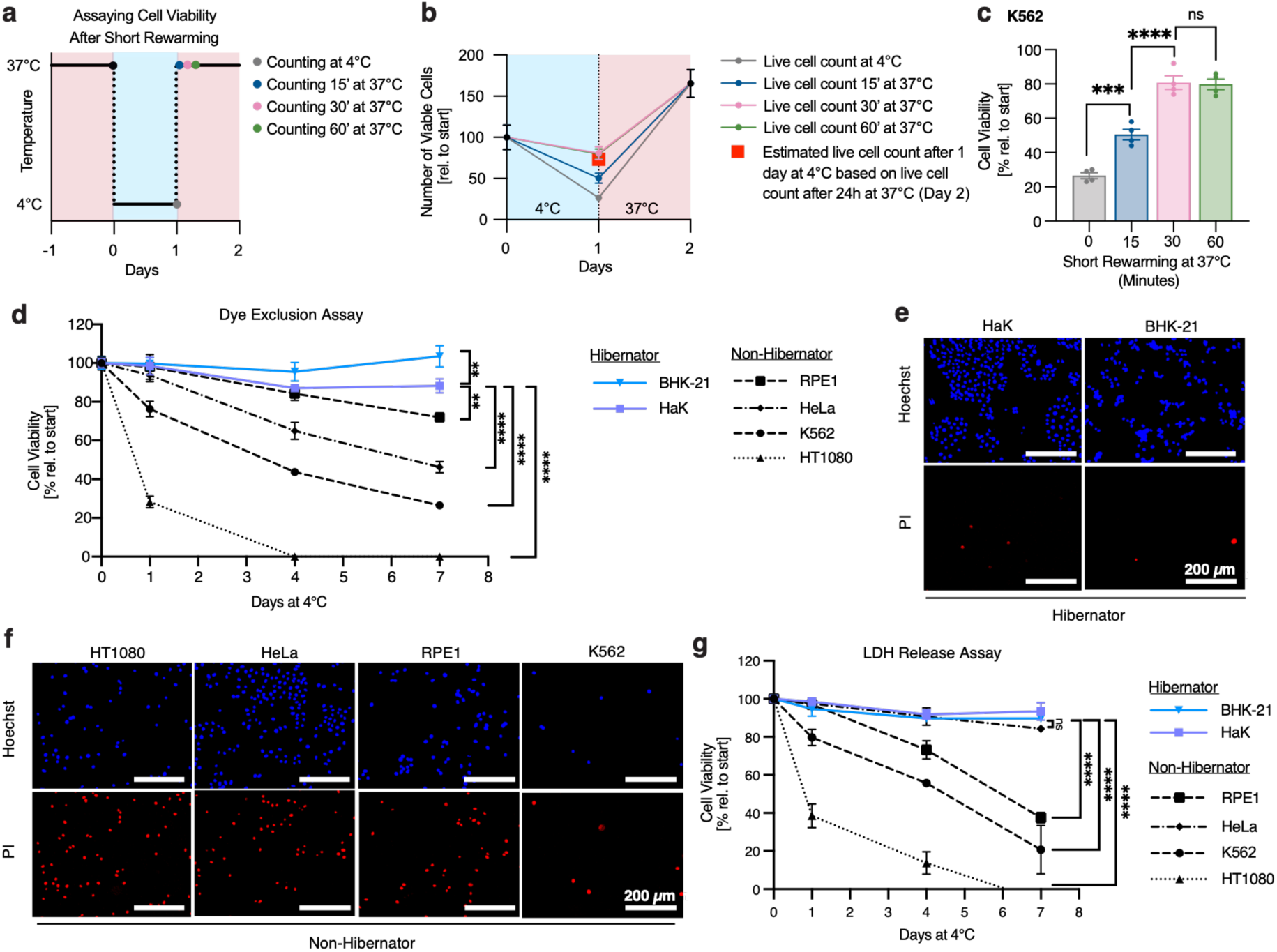
Hibernator-derived cells exhibit increased cold resistance. **a**, Schematic of cold cell viability counting, consisting of 1 day cold exposure (4°C), followed by a short rewarming at 37°C for 15, 30, or 60 minutes before trypan blue staining. **b,** Number of viable K562 cells based on trypan blue staining after one day at 4°C and subsequent rewarming for 24 hours at 37°C. Numbers are normalized to initial cell counts. Blue shaded regions indicate 4°C exposure and shaded pink regions indicate 37°C exposure. Red square indicates calculated cell counts after one day at 4°C based on the viable cell number measured after 24-hour rewarming. **c**, Viability of K562 cells was assessed by trypan blue staining after incubation at 37°C for 0, 15, 30, or 60 minutes following 24 hours at 4°C (*n* = 4). Cells incubated at 37°C for 15, 30, and 60 minutes show a significant increase in cell counts compared to cells counted without rewarming (n = 4 samples per condition, \*\*\**P* = 0.0007, \*\*\*\**P* < 0.0001). **d**, Viability of hibernator (BHK-21, HaK)- and non-hibernator (HeLa, RPE1, HT1080, K562)-derived cell lines at 4°C as measured by trypan blue staining. Hibernator-derived lines show significantly increased cold resistance compared to lines derived from non-hibernators at 7 days 4°C (*n* = 4 samples per data point, \*\*\*\**P* < 0.0001). **e, f**, Fluorescence images of hibernator- and non-hibernator-derived cell lines after 4 days at 4°C. Cultures were stained with 1 µg/mL Hoechst 33342 and 1 µg/mL propidium iodide (PI) to distinguish live and dead cells. **g**, Viability of hibernator- and non-hibernator-derived cell lines at 4°C as measured by LDH release (*n* = 4 samples per data point, \*\*\*\**P* < 0.0001). All values show mean ± SEM, with significance measured by one-way ANOVA adjusted for multiple comparisons by Dunnett’s test. \**P* < 0.05; \*\**P* < 0.01; \*\*\**P* < 0.001; \*\*\*\**P* < 0.0001; ns *P* > 0.05.

### Hibernator-derived cells show enhanced cold tolerance compared to human cells

Using our modified assay for cell viability after cold challenge, we examined the relative cold tolerance of hibernator and non-hibernator cells. We tested four commonly used human cell lines (HT1080, HeLa, RPE1, and K562), as well as two cell lines (BHK-21 and HaK) derived from Syrian hamsters (*Mesocricetus auratus*), a hibernating mammal. Cells were placed at 4°C - the lower end of the temperature range which Syrian hamsters reach during hibernation - and we assessed cell survival after 1, 4, and 7 days of cold exposure. Consistent with prior reports^5,7,8^, hamster-derived cell lines maintained high levels of viability, whereas human cell lines showed varying degrees of death (**Figures 1d-f**), with several exhibiting less than 50% survival following 7 days in the cold. Notably, we observed generally higher cell survival rates than previously reported^5^, likely owing to our modified method of measuring cell viability and culture conditions. Similar survival trends were also observed using an orthogonal assay based on lactate dehydrogenase (LDH) release from dead cells into the media (**Figure 1g**).

### Genome-scale CRISPR screen in a hibernator-derived cell line identifies suppressors of cold-induced cell death

To investigate the molecular pathways regulating cold resistance in hibernator-derived BHK-21 cells, we designed a genome-scale CRISPR-Cas9 screen. Taking advantage of a recent chromosome-level assembly of the *Mesocricetus auratus* genome^25^ and a modified CRISPOR-based guide selection algorithm (Methods), we generated a library of 218,143 single guide RNAs (sgRNAs) targeting all 21,473 annotated genes (∼10 guides per gene), including 2,299 intergenic-targeting and 250 non-targeting sgRNAs as negative controls. The pooled library was introduced into BHK-21 cells via a lentiviral vector expressing the sgRNA along with Cas9, achieving ∼1000-fold library representation with a multiplicity of infection (MOI) < 1.

Considering that many mammalian species, including Syrian hamsters, undergo prolonged periods of cold exposure (deep torpor) and rewarming (interbout arousals) during seasonal hibernation, we sought to recapitulate these temperature changes in our screen. We thus exposed BHK-21 cells to three cycles consisting of 4°C cold exposure (4 days) followed by rewarming at 37°C (2 days) (“4°C Cycling”, **Figure 2a**). To isolate genes selectively required upon cold exposure, we also analyzed BHK-21 cultures transduced in parallel that were maintained at 37°C for four passages/cycles (“37°C Control”, **Figure 2a**). For both paradigms, genomic DNA was isolated from cells during each cycle/passage to monitor selective sgRNA depletion. To assess our screen performance, we tested whether sgRNAs targeting previously identified human core essential genes^26^ were significantly depleted compared with nontargeting and intergenic sgRNAs and found that we can robustly distinguish between these functional categories at all three cycles and at both 37°C and 4°C (estimated p-value < 2.2e-16) (**Figure S3**).

**Figure 2.**
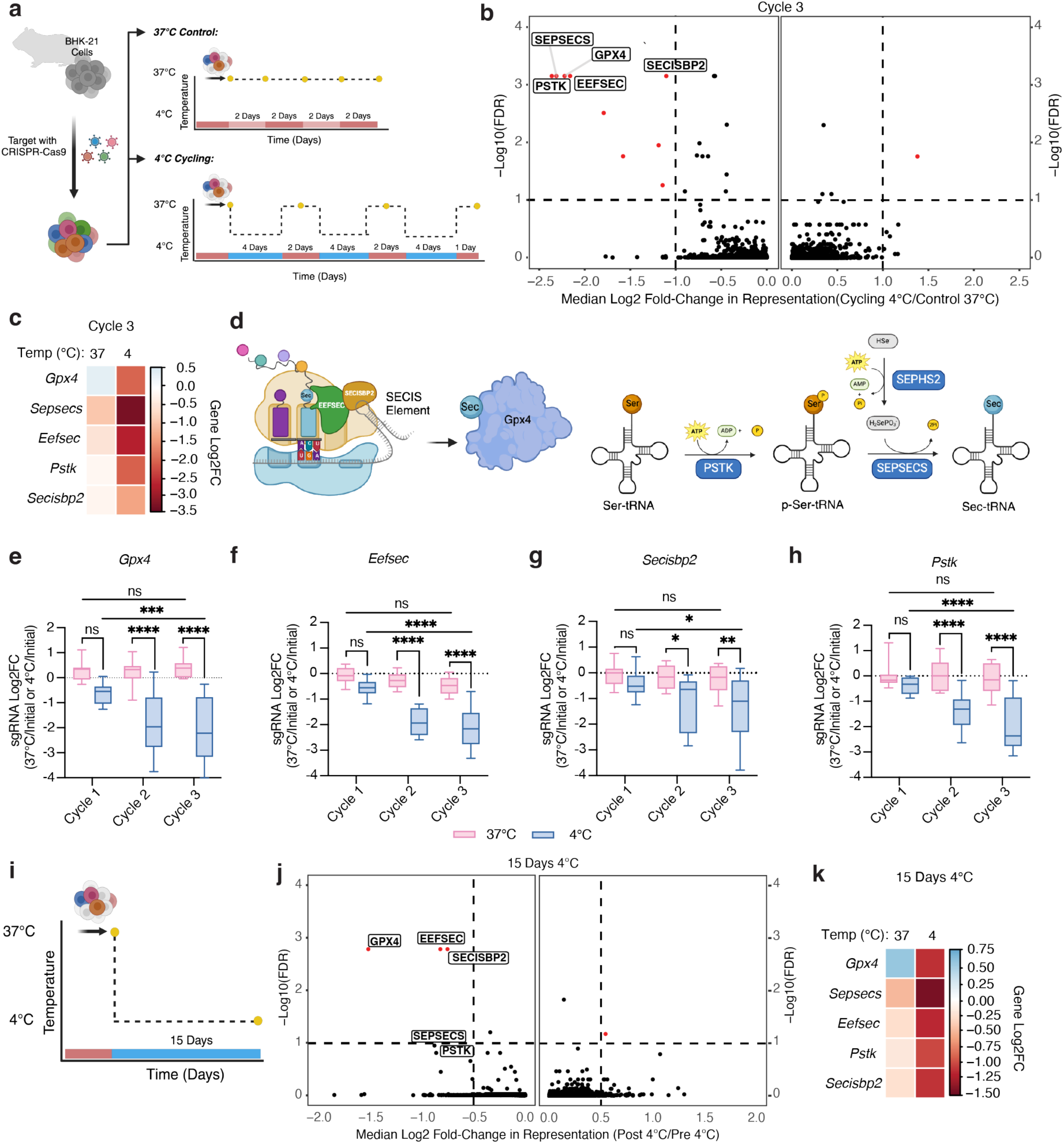
Unbiased CRISPR screens identify the GPX4 pathway as necessary for cold-induced survival in hibernator-derived BHK-21 cells. **a**, Schematic of CRISPR screen paradigm, consisting of four passages of control cells kept at 37°C and three cycles of cold exposure (4°C) for 4 days followed by rewarming (37°C) for 2 days. Yellow dots indicate points of sample collection. **b**, Volcano plot showing median log2 fold-change of genes after three cycles of cold exposure and rewarming (4°C) compared to cells following three passages at 37°C. Red dots indicate selectively required genes with a median log2 fold-change < -1 or > 1 and FDR < 0.1. **c**, Heatmap of the median Log2FC of ferroptosis-related genes after three cycles of cold exposure and rewarming (4°C) compared to three passages at 37°C. **d**, Partial schematic of the selenocysteine incorporation pathway. Left: Production of the GPX4 selenoprotein requires recoding of a UGA codon to the amino acid selenocysteine (Sec). This process involves a cis-acting SECIS element within the *Gpx4* mRNA 3’ UTR, SECIS binding protein 2 (SECISBP2), a specific eukaryotic elongation factor (EEFSEC), and Sec-charged tRNA. Right: The Sec-charged tRNA is generated by the combined action of Phosphoseryl-tRNA kinase (PSTK), Selenophosphate synthetase 2 (SEPHS2), and selenocysteine synthase (SEPSECS). **e-h**, Median log2 fold-change (Log2FC) of 10 guides per targeted gene, showing guide depletion over three cycles of cold exposure and rewarming. Significance between Cycle 1 versus Cycle 3 is measured by two-way ANOVA adjusted for multiple comparisons by Dunnett’s test. Significance between 37°C and 4°C for each cycle is measured by two-way ANOVA adjusted for multiple comparisons by Bonferroni’s test. **e**, *Gpx4*, **f**, *Eefsec*, **g**, *Secisbp2,* **h**, *Pstk*. **i**, Schematic of CRISPR screen paradigm, showing cells exposed to 4°C continuously for 15 days. Yellow dot indicates point of sample collection. **j**, Volcano plot showing median log2 fold-change in abundance of guides targeting the indicated genes after 15 days of 4°C exposure compared to one passage at 37°C. Red dots indicate selectively required genes with a median log2 fold-change < -0.5 or > 0.5 and FDR < 0.1. **k**, Heatmap of the median Log2FC in abundance of guides targeting ferroptosis-related genes after 15 days of 4°C exposure compared to 37°C control cultures. \**P* < 0.05; \*\**P* < 0.01; \*\*\**P* < 0.001; \*\*\*\**P* < 0.0001; ns *P* > 0.05.

To identify genes that selectively modify (suppress or potentiate) cold-induced cell loss, we compared the guides depleted during three cycles of cold exposure and rewarming to controls at 37°C with matched population doublings. Among the 21,473 targeted genes, only nine genes appeared selectively required during cold exposure (FDR < 0.1, log2 fold-change [Log2FC] >1, **Figure 2b, Table S1, S2**). Four selectively required genes - *Dld*, *Lias*, *Lipt1*, and *Ybey –* are involved in lipoylation and mitochondrial RNA processing, suggesting that changes to mitochondrial metabolism sensitize cells to cold exposure or rewarming^27–31^.

Strikingly, five of the nine required genes - *Gpx4*, *Eefsec*, *Pstk*, *Secisbp2*, and *Sepsecs -* represent known components of the glutathione/Glutathione Peroxidase 4 (GPX4) antioxidant pathway, indicating that this pathway functions as a potent suppressor of cold-induced cell death in hibernator-derived BHK-21 cells (**Figure 2c**). *Gpx4* encodes a selenocysteine-containing lipid antioxidant enzyme that acts to suppress lipid peroxidation via the glutathione-mediated reduction of lipid hydroperoxides to non-toxic lipid alcohols^32,33^. GPX4 inhibition sensitizes cells to ferroptosis, a distinct form of programmed cell death characterized by iron-dependent accumulation of lipid peroxidation^34–36^. Notably, cold exposure has previously been associated with increased lipid peroxidation and ferroptosis in human cells^5^. Indeed, we also observed an increase in the oxidation of the lipophilic dye, C11 BODIPY in K562, RPE1, and BHK-21 cells exposed to 4°C for 24 hours (**Figure S4**), consistent with increased lipid peroxidation. Underscoring the requirement for Gpx4 activity, four of the eight remaining genes selectively required upon cold exposure (*Eefsec, Secisbp2, Pstk, Sepsecs*) are directly or indirectly involved in selenocysteine incorporation into cellular proteins and thus required for Gpx4 activity (**Figures 2d-h**)^37–44^. Given that our initial screen design involved repeated rewarming cycles, we could not exclude the possibility that the Gpx4 pathway confers resistance to rewarming-associated stress rather than cold tolerance per se. To address this issue, we took advantage of the high cold tolerance of BHK-21 cells and included a second screen arm using similar methods in which cells were continuously exposed to 4°C for 15 days (**Figure 2i**). Sequencing prior to and following 15 days of cold exposure, followed by a brief rewarming to 37°C, confirmed significant depletion (FDR < 0.1, Log2FC < -0.5) of sgRNAs targeting *Gpx4*, *Eefsec*, and *Secisbp2* (as well as a trend toward depletion for *Sepsecs* and *Pstk*), confirming a critical role for the Gpx4 pathway in BHK-21 cell prolonged cold tolerance (**Figures 2j,k, Table S2, S3**). Thus, two unbiased genome-scale screen arms identified Gpx4 as a major suppressor of cold-induced cell death in hibernator-derived BHK-21 cells in the context of both cyclical and continuous cold exposure.

### Gpx4 catalytic activity is essential for hibernator cell survival during cold exposure

To further validate the role of GPX4 in cold-mediated cell death, we employed the top-ranked sgRNA from our screen to generate clonal *Gpx4* knockout BHK-21 cells. We sorted individual cells to obtain clonal knockout cell lines and maintained them in the presence of liproxstatin-1 since they showed stunted growth at 37°C (**Figure S5**). We confirmed the absence of Gpx4 via immunoblotting (**Figure 3a**). While wild-type (WT) BHK-21 cells maintained high cell viability at 4°C, all three *Gpx4* knockout lines exhibited complete cell death within 4 days of cold exposure (\*\*\*\**P* < 0.0001) (**Figure 3a)**. Moreover, these differences reflected Gpx4 loss, as opposed to off-target effects insofar as lentiviral re-expression of cytosolic hamster Gpx4 (\*\*\*\**P* < 0.0001), but not a GFP control gene or a catalytically dead mutant form of Gpx4 (mGPX4) that has a serine in the active site instead of a selenocysteine, restored cold tolerance of *Gpx4* knockout lines (**Figures 3b,c).**

**Figure 3.**
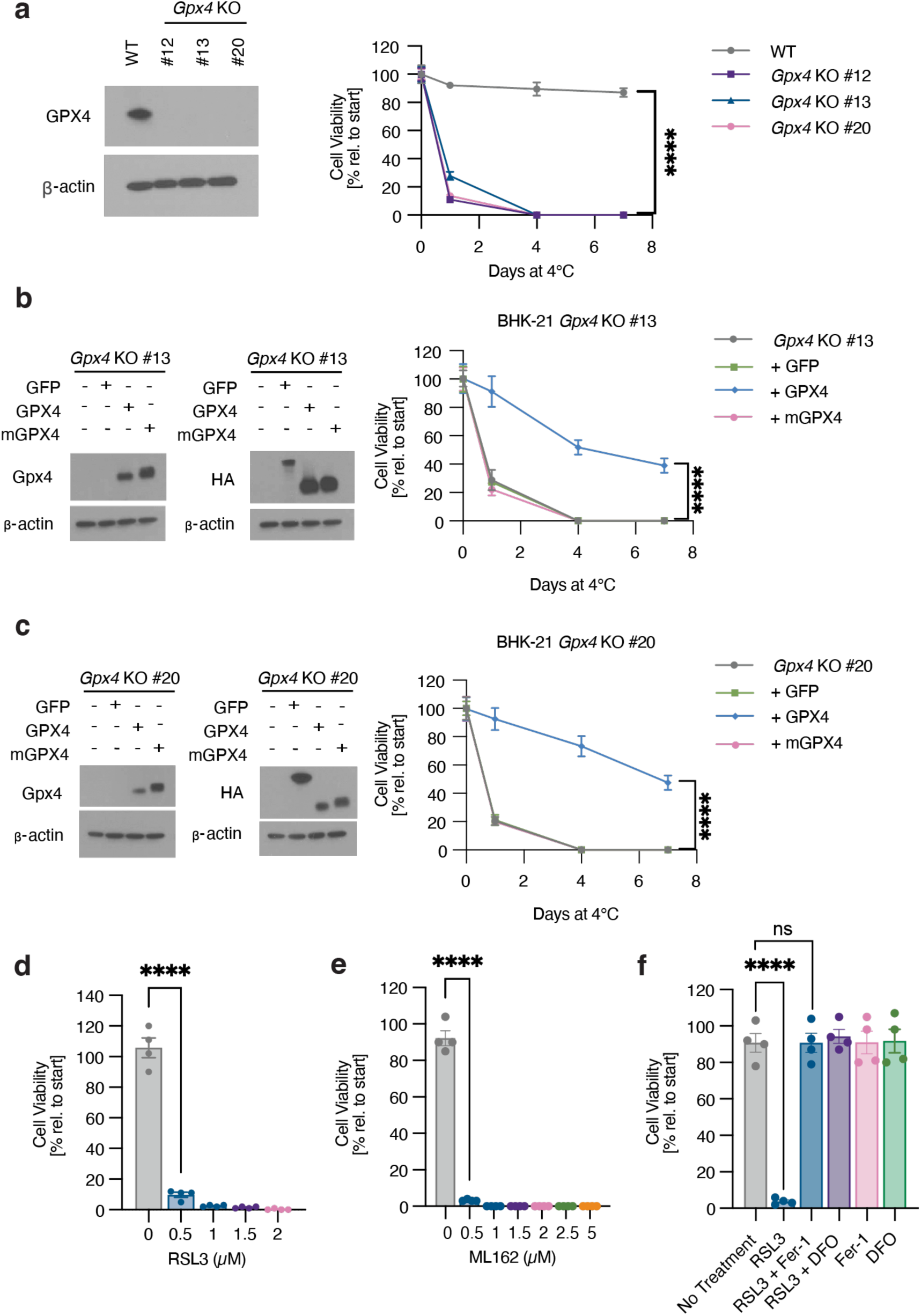
GPX4 activity is required for BHK-21 cell cold tolerance. **a**, Stable *Gpx4* knockout (KO) BHK-21 cell lines exhibit reduced cold tolerance. Left: Western blot of wild-type (WT) and individual *Gpx4* KO clone lysates for GPX4 and β-actin loading control. Right: Viability of *Gpx4* KO lines is significantly lower than WT BHK-21 cells at 7 days 4°C by trypan blue staining (*n* = 4, \*\*\*\**P* < 0.0001), with complete cell death by four days at 4°C. **b, c**, Reintroduction of wild-type Syrian hamster GPX4 (GPX4), but not a catalytically dead form of GPX4 (mGPX4) rescues cold-induced cell death in two independent BHK-21 *Gpx4* KO clonal cell lines. Left panels: Western blots for HA and GPX4 along with β-actin loading control. Right panels: Expression of WT hamster GPX4 showed significantly higher cell viability at 7 days 4°C compared to the corresponding parental *Gpx4* KO, GFP-, and mGPX4-expressing lines by trypan blue staining (*n* = 4, \*\*\*\**P* < 0.0001). **d, e**, Treatment with the GPX4 inhibitors RSL3 (**d**) or ML162 (**e**) results in enhanced cold-induced death in BHK-21 cells by 4 days at 4°C as measured by trypan blue exclusion (*n* = 4, \*\*\*\**P* < 0.0001). **f**, Cold-induced BHK-21 cell death upon RSL3 treatment occurs via ferroptosis. BHK-21 cells were placed at 4°C and treated with RSL3 (1 µM) and/or the ferroptosis inhibitor ferrostatin-1 (Fer-1, 1 µM) or iron chelator DFO (100 µM) for 4 days (*n* = 4). Treatment with RSL3 resulted in significantly lower cell viability than no treatment as determined by one-way ANOVA adjusted for multiple comparisons by Tukey’s HSD (\*\*\*\**P* < 0.0001). All values show mean ± SEM, with significance measured by two-tailed t test, unless otherwise indicated. \**P* < 0.05; \*\**P* < 0.01; \*\*\**P* < 0.001; \*\*\*\**P* < 0.0001; ns *P* > 0.05.

Together, these genetic experiments strongly implicate the catalytic activity of Gpx4 in BHK-21 cold tolerance. However, the constitutive nature of these interventions precluded a determination of whether Gpx4 functions to actively oppose cell death during cold-exposure or whether loss of its catalytic activity prior to cold exposure sensitizes cells to subsequent cold challenge. To address this issue, we used RSL3 and ML162, two competitive small-molecule Gpx4 inhibitors^45–47^. Acute treatment of cultured BHK-21 cells with these inhibitors during cold exposure yielded dose-dependent increases in cell death. Indeed, by four days of treatment, both RSL3- and ML162-treated BHK-21 cells display significantly increased cold-induced death compared to untreated controls (\*\*\*\**P* < 0.0001) (**Figures 3d,e**). Importantly, relative to WT cells, *Gpx4* knockout cells exhibited no significant increase in cold-induced cell death upon RSL3 treatment following a short cold exposure (8 hours) (**Figure S6**), confirming the specificity of these inhibitors under the test conditions. We therefore conclude that Gpx4 catalytic activity is required in BHK-21 cells during the course of cold exposure to actively oppose cold-induced cell death.

Given that Gpx4 activity acts as a major endogenous suppressor of ferroptosis^34^, we tested whether the cold-induced cell death observed under Gpx4 inhibition occurred via the ferroptosis pathway. To this end, we made use of two small molecule inhibitors of ferroptosis, ferrostatin-1, a lipophilic antioxidant, and deferoxamine mesylate (DFO), a Fe^2+^ chelator^48,49^. Both inhibitors effectively rescued cold-induced cell death in the presence of the Gpx4 inhibitor RSL3^50^ (**Figure 3f**), indicating that acute inhibition of Gpx4 activity in the cold drives cell death via ferroptosis.

Together, through a combination of genetic and pharmacological approaches, our studies demonstrate that Gpx4 activity confers substantial cold resistance in hibernator-derived cells, protecting cells from cold-induced ferroptosis.

### Genome-scale CRISPR screen identifies determinants of cold sensitivity in human cells

In contrast to hibernator-derived cell lines, human cells display varying cold sensitivities in culture, with some exhibiting pronounced intolerance to the cold (**Figure 1d-g**). Consistent with prior reports^5,8,51^, pharmacological experiments in several human cell lines (K562, HT1080, RPE1, and HeLa cells) confirmed that cold-induced cell-loss occurred primarily via ferroptosis, with both antioxidant ferroptosis inhibitors (ferrostatin-1 and liproxstatin-1) and iron chelators (DFO and 2’2’-pyridine) increasing the cold survival in all four cell lines in a dose-dependent manner. By contrast, neither the Caspase inhibitor Z-VAD-FMK nor the necroptosis inhibitor necrostatin-1 significantly affected cell viability (**Figure S7, S8**).

To gain further insight into the genetic modifiers of human cell cold sensitivity and resistance, we designed and performed an additional CRISPR-Cas9 screen in cold-sensitive human K562 cells exposed to cold temperatures. A pooled lentiviral CRISPR-Cas9 library targeting 19,381 genes (∼5 sgRNAs per gene) was delivered to K562 cells at an MOI of ∼0.5. Transduced cells were placed at 4°C for five days to allow for cold-induced cell death to occur in approximately 79 ± 1.76% of cells, followed by a three-day rewarming to 37°C to allow for amplification of remaining cells (**Figure 4a**). This cold-rewarming cycle was repeated three times, with cells collected after each cycle to monitor the relative enrichment and depletion of individual sgRNAs. As in our prior screens, separate cultures transduced in parallel were maintained at 37°C for comparative analysis. In addition, to discriminate between ferroptosis-dependent and - independent regulators of cold tolerance, we conducted a parallel screen in cold-exposed K562 cells under continuous ferrostatin-1 treatment. As before, the efficiency and specificity of the screens were assessed based on effective depletion of characterized core essential genes^26^ as well as the lack of depletion of negative control guides (250 nontargeting and 125 intergenic sgRNAs) (**Figure S9)**.

**Figure 4.**
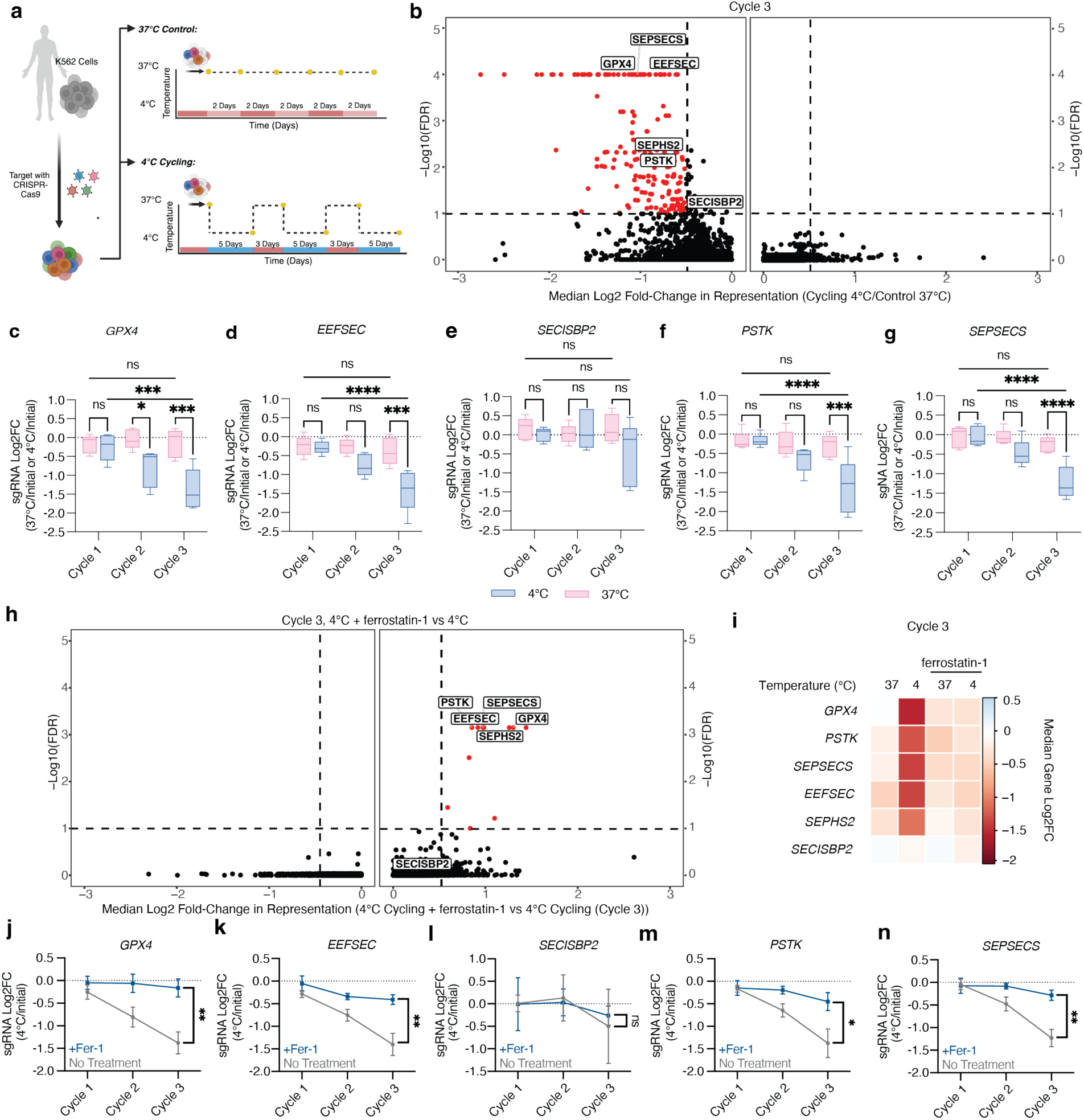
Genome-wide CRISPR screens identify GPX4 as a suppressor of cold-induced cell death in human cells. **a**, Schematic of CRISPR screen paradigm, consisting of three cycles of 5 days of cold exposure (4°C) interrupted by 3-day rewarming (37°C) periods. Yellow dots indicate points of sample collection. **b**, Volcano plot showing median log2 fold-change in abundance of guides targeting the indicated genes after three cycles of 4°C cold exposure compared to three passages at 37°C (Control). Red dots indicate selectively required genes with a median log2 fold-change < - 0.5 or > 0.5 and FDR < 0.1. **c-g**, Combined median log2 fold-change (Log2FC) of 5 sgRNAs per targeted gene, showing sgRNA depletion over three cycles of cold exposure. Significance between Cycle 1 versus Cycle 3 is measured by two-way ANOVA adjusted for multiple comparisons by Dunnett’s test. Significance between 37°C and 4°C for each cycle is measured by two-way ANOVA adjusted for multiple comparisons by Bonferroni’s test. **c**, *GPX4*, **d**, *EEFSEC*, **e**, *SECISBP2*, **f**, *PSTK*, **g**, *SEPSECS*. **h**, Volcano plot showing median log2 fold-change in abundance of guides targeting the indicated genes after three cycles of 4°C cold exposure in the presence or absence of 1 µM of ferrostatin-1. Red dots indicate selectively required genes with a log2 fold-change < -0.5 or > 0.5 and FDR < 0.1. **i**, Heatmap of the median Log2FC in abundance of guides targeting ferroptosis-related genes after three cycles of cold exposure (4°C) compared to 37°C control cultures with and without ferrostatin-1. **j-n**, Depletion of 5 sgRNAs per gene over three cycles of cold exposure and rewarming with and without ferrostatin-1 (1 uM) treatment. **j**, *GPX4* (\*\**P* = 0.0047), **k**, *EEFSEC* (\*\**P* = 0.0059), **l**, *SECISBP2* (ns, *P* = 0.5457), **m**, *PSTK* (\**P* = 0.0393), **n**, *SEPSECS* (\*\**P* = 0.0029) as measured by two-tailed t-test at Cycle 3. All values show mean ± SEM. \**P* < 0.05; \*\**P* < 0.01; \*\*\**P* < 0.001; \*\*\*\**P* < 0.0001; ns *P* > 0.05.

To our surprise, this screen did not identify any genes whose disruption significantly enhanced K562 cold tolerance. By contrast, we identified 176 genes selectively required during cold cycling relative to constant 37°C control conditions (FDR < 0.1, Log2FC < -0.1, and requiring >2 sgRNAs, **Figure 4b**). It is notable that K562 cold survival was sensitive to the targeting of larger number of genes than the BHK-21 line, potentially indicating increased dependencies on genetic pathways which are not essential in BHK-21 cells (**Figure S10, Table S6**). Among the pathways identified, we honed in on the selenocysteine incorporation pathway and observed *GPX4* and the selenocysteine incorporation genes *PSTK*, *SEPSECS*, and *EEFSEC* within the top-ranked K562 cold-protective genes, suggesting that the GPX4 pathway confers significant resistance to the cold not only in cold-tolerant hibernator cells, but also in cold-sensitive human cells (**Figures 4c-g**). Notably, we identified 11 genes that were no longer essential (Fold enrichment > 0.50, FDR < 0.10) in the presence of Ferrostatin-1, indicating that these genes act to suppress cold-induced ferroptosis (**Figures 4h-n, Table S4, S5**). Although multiple pathways have been described as suppressors of ferroptosis in other contexts^52–56^, it is notable that 5 of 11 identified genes represent known components of the GPX4 pathway.

To confirm these findings, we generated three clonal *GPX4* knockout K562 cell lines and confirmed the loss of GPX4 via immunoblotting (**Figure 5a**). Similar to our findings in BHK-21 cells, K562 *GPX4* knockout clones exhibited stunted growth and were expanded in the presence of liproxstatin-1 (**Figure S5b**). Importantly, GPX4 loss greatly increased cold-induced cell death (\*\*\*\**P* < 0.0001) (**Figures 5a**), thus confirming an essential role for *GPX4* in human cell cold tolerance. Similar effects were also observed upon acute pharmacological inhibition of GPX4 with RSL3 or ML162 at 4°C (\*\*\*\**P* < 0.0001) (**Figures 5b, c**). These effects were dependent on GPX4 inhibition insofar as the cold tolerance of K562 *GPX4* knockout cells was unaffected by RSL3 treatment (**Figure S11**). In addition, RSL3-driven cold-induced cell death was rescued by either ferrostatin-1 or DFO treatment, indicating that RSL3-induced cell loss occurs via the ferroptosis pathway (**Figure 5d**). Increased cold-induced death in the *GPX4* knockout K562 cell lines could be rescued upon lentiviral expression of either the cytosolic human (hs) or hamster (ma) GPX4, but not GFP or catalytically dead GPX4 variants (**Figure 5e).** Together, our genetic and pharmacological studies reveal that K562 cells, despite being significantly less cold-tolerant than hamster BHK-21 cells, also rely on the GPX4 pathway to suppress cold-induced ferroptosis.

**Figure 5.**
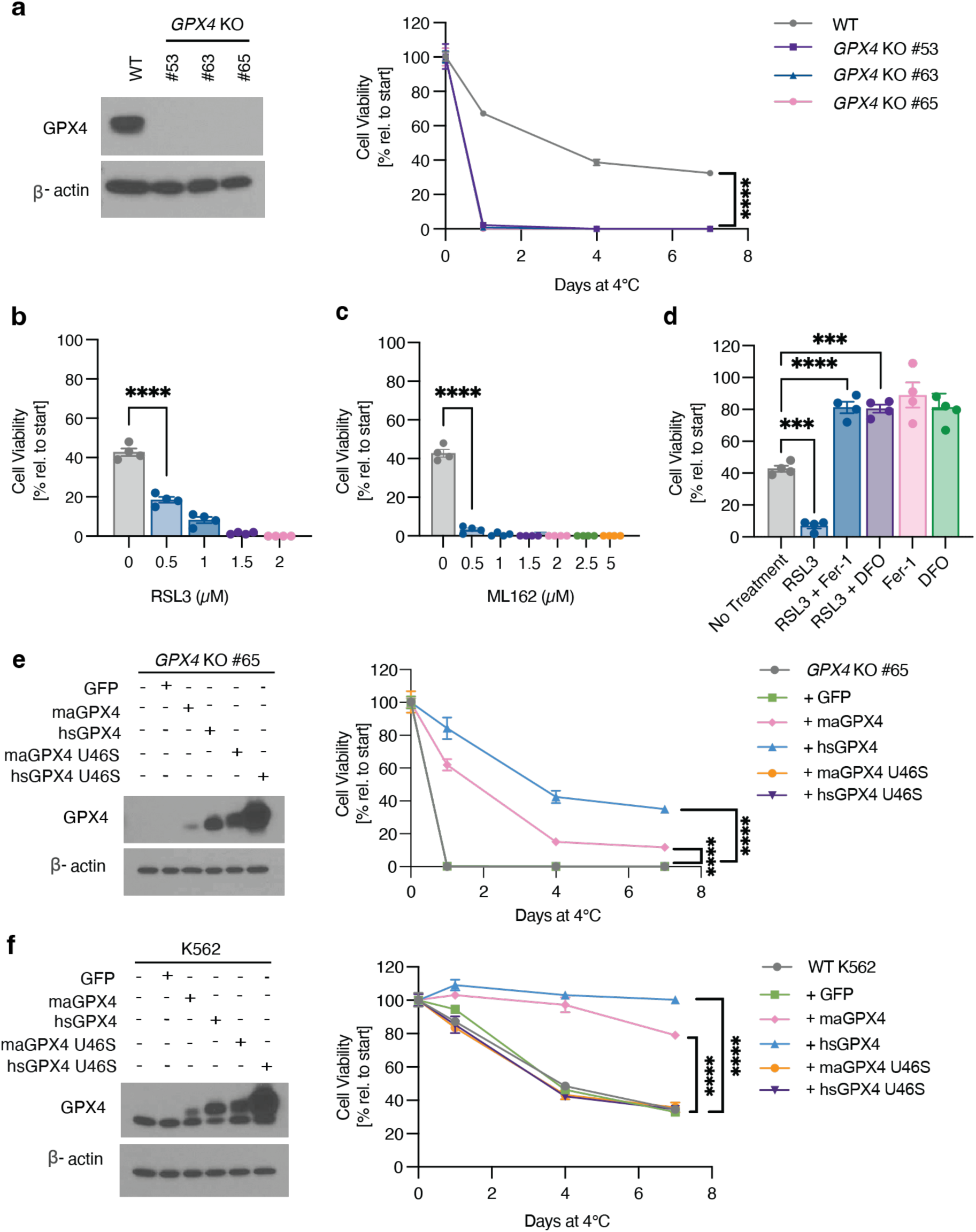
Endogenous GPX4 activity is limiting for K562 cell cold tolerance. **a**, Stable *GPX4* knockout (KO) K562 cell lines exhibit reduced cold tolerance. Left: Western blotting of wild-type (WT) and individual *GPX4* KO clones for GPX4 and β-actin loading control. Right: Viability of *GPX4* KO lines is significantly lower than WT K562 cells as measured by trypan blue staining (*n* = 4, \*\*\*\**P* < 0.0001), with complete cell death by one day at 4°C. Significance measured by two-tail t-test at 7 days 4°C. **b, c**, Treatment with the GPX4 inhibitors RSL3 (**b**) or ML162 (**c**) results in enhanced K562 cold-induced death by 4 days at 4°C (*n* = 4, \*\*\*\**P* < 0.0001) as measured by two-tailed t-test. **d**, Cold- and RSL3-induced K562 cell death occurs via ferroptosis. K562 cells were placed at 4°C and treated with RSL3 (1 µM) and/or the ferroptosis inhibitor ferrostatin-1 (Fer-1, 1 µM) or iron chelator DFO (5 µM) for 4 days (*n* = 4). Treatment with RSL3 resulted in significantly lower cell viability than no treatment as determined by one-way ANOVA adjusted for multiple comparisons by Tukey’s HSD (\*\*\**P* = 0.0002). **e**, Reintroduction of wild-type human (hs) or hamster (ma) GPX4, but not catalytically dead forms of GPX4 (Gpx4 U46S), rescues cold-induced cell death in a K562 *GPX4* KO clonal cell line (*n* = 4). Left: Western blot for GPX4 levels and β-actin loading control. Right: Expression of WT human GPX4 and hamster GPX4 resulted in significantly higher cell viability compared to the corresponding parental *GPX4* KO, GFP-, and mutGPX4-expressing lines by trypan blue staining, as measured by one-way ANOVA adjusted for multiple comparisons by Dunnett’s test at 7 days 4°C (\*\*\*\**P* < 0.0001). **f**, Overexpression of wild-type human or hamster GPX4, but not catalytically dead forms of GPX4 (mutGPX4), suppresses cold-induced cell death in a K562 cells (n = 4). Left: Western blot for GPX4, with β-actin loading control. Right: Expression of WT human GPX4 and hamster GPX4 resulted in significantly higher cell viability compared to wild-type, GFP-, and mutGPX4-expressing K562 lines by trypan blue staining, as measured by one-way ANOVA adjusted for multiple comparisons by Dunnett’s test at 7 days 4°C (\*\*\*\**P* < 0.0001). All values show mean ± SEM. \**P* < 0.05; \*\**P* < 0.01; \*\*\**P* < 0.001; \*\*\*\**P* < 0.0001; ns *P* > 0.05.

### GPX4 catalytic activity is limiting for K562 cell cold tolerance

Given that the GPX4 pathway is active during cold exposure and essential for cold survival in both hamster BHK-21 and human K562 cells, it is somewhat surprising that these cell lines exhibit markedly different cold tolerances. In this regard, it has previously been suggested^5^ that upon cold exposure human cells, in contrast to hibernator-derived cells, rapidly deplete intracellular glutathione, an essential co-factor for GPX4 function, which could thus account for their increased cold sensitivity. To test this idea, we supplemented K562 growth media with either cell-permeable Glutathione reduced ethyl ester (GSH-MEE), N-acetyl-cysteine (NAC), or L-Cystine, a glutathione precursor; however, these interventions failed to significantly increase K562 cells’ cold tolerance under our culture conditions (**Figure S12**). We therefore examined the possibility that GPX4 protein levels in K562 cells are limiting for their cold tolerance. To this end, we generated K562 lines overexpressing either human or hamster GPX4. Both of these overexpression lines exhibited strikingly improved cold tolerance compared to the wild-type K562 parental line and a GFP-overexpressing control cell line (\*\*\*\**P* < 0.0001) (**Figure 5f)**.

Indeed, GPX4-overexpressing K562 cells displayed comparable cold tolerance to that of hibernator-derived BHK-21 cells (**Figure 1d, g**). To confirm that increased GPX4 catalytic activity was responsible for the improved cell cold tolerance, we generated additional K562 lines overexpressing catalytically dead forms of hamster or human GPX4 in which the active site selenocysteine was mutated to serine and found that these GPX4 mutant lines exhibited no increase in cold tolerance (**Figure 5f**). Taken together, these results strongly suggest that GPX4 abundance serve as a key limiting determinant of K562 cell cold tolerance. Moreover, our finding that human and hamster GPX4 overexpression confer comparable increases in K562 cold tolerance may indicate that the intrinsic activity of the hamster and human GPX4 enzymes are similar.

### GPX4 activity is broadly required for cold tolerance in primary cells across evolutionarily distant mammalian species

Given that our studies had largely employed a small number of transformed cell lines, we sought to extend our investigation to examine cold tolerance in primary cell types as well as cells derived from additional hibernating species. To this end, we obtained primary and immortalized fibroblasts from human, mouse, and rat as well as from Syrian hamster, 13-lined ground squirrel (*Ictidomys tridecemlineatus*), and horseshoe bat (*Rhinolophus ferrumequinum*), three distantly related mammalian hibernators, and characterized their cold tolerance. While hibernator-derived fibroblasts exhibited robust, roughly uniform cold tolerance, we observed a surprisingly variable degree of cold tolerance across non-hibernator primary cells (**Figure 6a**), with human dermal fibroblasts exhibiting a remarkable 79 ± 4.93% cell viability after seven days at 4°C, comparable to that observed with hamster-derived BHK-21 cells.

**Figure 6.**
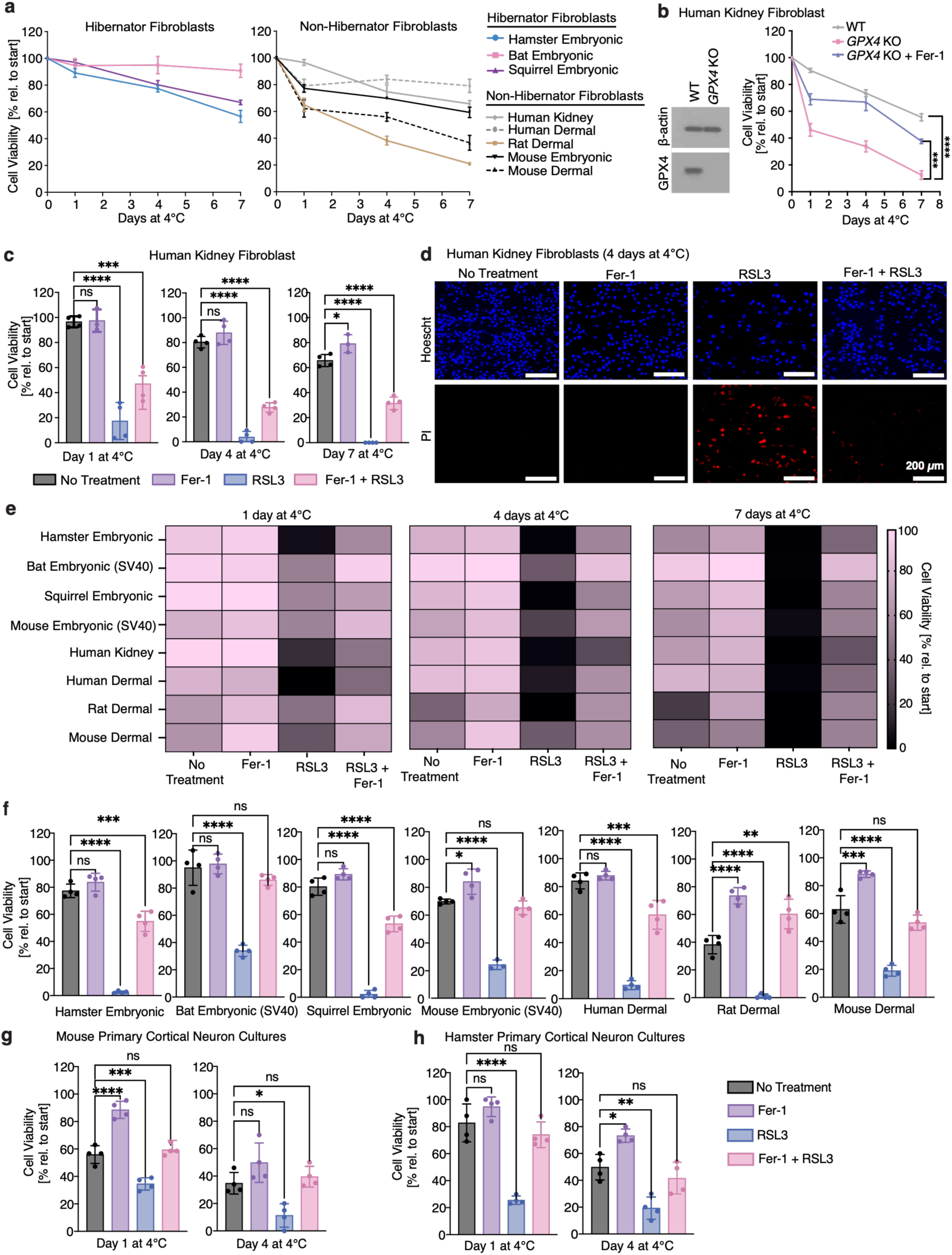
GPX4 is required for cold tolerance across both several hibernator and non-hibernator mammalian species and cell types. **a**, Viability of hibernator cells (Syrian hamster embryonic fibroblasts, Greater horseshoe bat embryonic fibroblasts, and 13-lined ground squirrel embryonic fibroblasts) and non-hibernator cells (SV40-immortalized mouse embryonic fibroblasts, human adult kidney fibroblasts, human adult dermal fibroblasts, rat adult dermal fibroblasts, and mouse adult dermal fibroblasts) exposed to 4°C as measured by trypan blue staining (n = 4, \*\*\*\**P* < 0.0001). **b**, Human kidney fibroblast *GPX4* knockout cells exhibit reduced cold tolerance compared to WT cells. Left: Western blot of wild-type (WT) and *GPX4* KO cells for GPX4 and β-actin loading control. Right: Viability of *GPX4* KO cells is significantly lower than WT cells at 4°C as measured by trypan blue staining (n = 4, \*\*\*\**P* < 0.0001). **c**, Gpx4 activity is essential for cold survival of primary human kidney fibroblasts. Human kidney fibroblasts were placed at 4°C and left untreated or treated with RSL3 (1 μM) and/or the ferroptosis inhibitor ferrostatin-1 (Fer-1, 1 μM) for 7 days (n = 4, \*\*\*\**P* < 0.0001). **d**, Representative fluorescence images of human kidney fibroblasts after 4 days at 4°C with no treatment, 1 μM Fer-1, 1 μM RSL3, or 1 μM Fer-1 and 1 μM RSL3. Cultures were stained with Hoechst 33342 and propidium iodide (PI) to identify live cells. **e**, Gpx4 activity is essential for cold survival in hibernator cells (Syrian hamster embryonic fibroblasts, Greater horseshoe bat embryonic fibroblasts, and 13-lined ground squirrel embryonic fibroblasts) and non-hibernator cells (SV40-immortalized mouse embryonic fibroblasts, human adult kidney fibroblasts, human adult dermal fibroblasts, rat adult dermal fibroblasts, and mouse adult dermal fibroblasts). Cells were placed at 4°C and left untreated, treated with RSL3 (1 μM), and/or ferrostatin-1 (Fer-1, 1 μM) for 7 days (n = 4). **f**, Expanded Day 4 data from **e)** indicates that Gpx4 activity is essential for fibroblast survival in the cold across several hibernator and non-hibernator species. Cells were placed at 4°C and left untreated or treated with RSL3 (1 μM) and/or ferrostatin-1 (Fer-1, 1 μM) for 7 days (n = 4, \*\*\*\**P* < 0.0001). **g-h**, Gpx4 activity is essential in **g)** mouse primary cortical neuron cultures and **h)** hamster primary cortical neuron cultures. Cells were placed at 4°C and left untreated, treated with RSL3 (1 μM) and/or the ferroptosis inhibitor ferrostatin-1 (Fer-1, 1 μM) for 1 or 4 days (n = 4). RSL3 treatment increased cell death, which was rescued by ferrostatin-1. All values show mean ± SEM, with significance measured by one-way ANOVA adjusted for multiple comparisons with Tukey’s HSD. \**P* < 0.05; \*\**P* < 0.01; \*\*\**P* < 0.001; \*\*\*\**P* < 0.0001; ns *P* > 0.05.

To examine the role of GPX4 in primary cell cold resistance, we transduced cells with Cas9 and sgRNAs targeting human *GPX4* to create a population of *GPX4* knockout human kidney fibroblasts (**Figure 6b**). Consistent with our prior findings in cell lines, GPX4 loss resulted in significantly decreased viability in the cold, which was largely rescued by ferrostatin-1 treatment in fibroblast cells (**Figure 6b**). Indeed, RSL3 treatment significantly decreased the viability of wild-type cold-exposed human kidney fibroblasts **(Figures 6c, d)**, and these effects were dependent on GPX4 inhibition as the cold tolerance of human kidney fibroblast *GPX4* knockout cells was largely unaffected by RSL3 treatment (**Figure S5c**).

To more widely examine the dependence of primary cell cold tolerance on GPX4 activity, we obtained and tested fibroblasts derived from six mammalian species. We found the GPX4 activity broadly contributes to cold tolerance across hibernators and non-hibernators, as RSL3-treated cells showed increased cold-induced cell death that was largely rescued by ferrostatin-1 treatment (**Figures 6e, f**). We also extended these findings to a non-fibroblast cell type, obtaining similar results with primary cortical neuronal cultures prepared from neonatal mice and hamsters (**Figures 6g, h**). Taken together, our data indicate that primary cell types are strongly sensitive to GPX4 loss in the cold and suggest that their differential cold sensitivities may reflect different levels of GPX4-mediated cold tolerance.

## Discussion

Hibernators can lower their body temperature to ∼4°C for several days to weeks, indicating that their cells and tissues possess the ability to survive extended periods of cold. Previous studies have indicated that ferroptosis contributes to cold-induced cell death in cultured human cells^5,8,51^ and identified several genes that modulate cold tolerance^57,58^. However, the genetic modifiers of cold sensitivity in hibernators and non-hibernators have yet to be systematically explored. Here, we conducted a series of unbiased genome-wide CRISPR-Cas9 screens in both hibernator- and non-hibernator-derived cells to investigate the mechanisms controlling cellular cold tolerance.

Our findings, further validated using stable genetic knockout lines and pharmacological inhibitors, identify the GPX4 pathway as a critical suppressor of cold-induced cell death. Indeed, overexpression studies in human K562 cells suggest that GPX4 abundance can serve as a key limiting determinant of cellular cold tolerance. Notably, Sone et al.^57^ recently also identified GPX4 as a strong regulator of cold tolerance through overexpression screening. These independent findings further support the current study identifying GPX4 as a conserved suppressor of cold-induced mammalian cell death. The consistency of our observations across diverse cell lines and primary cells, including cells derived from six different mammalian species, argues that GPX4 may serve as an essential and evolutionarily conserved mechanism to protect cells from cold-induced ferroptotic cell death.

During the course of our studies, we used trypan blue, a commonly employed membrane-impermeable dye, to assess cell viability. We made the surprising observation that cells stained with trypan blue or TO-PRO-3 (a fluorescent membrane impermeable dye) exhibit increased dye retention immediately after cold exposure, an artifact not reflecting genuine cell death.

Rather, cell counting following brief rewarming yields more accurate results, in some cases reflecting significantly higher cell viability than previously appreciated. Employing this modified approach to measure cold-induced cell death across sixteen cell lines and primary cell types points to more nuanced variations in cold survival across cell types. These differences are highlighted by the pronounced contrast between the poor cold viability of human HT1080 cells and the robust cold tolerance observed in human fibroblasts. The underlying basis of this transient cell permeability for trypan blue following cold exposure remains unclear; however, it is noteworthy that in *S. cerevisiae* heat and chemical insults have been found to induce a brief window of membrane permeability to PI prior to membrane repair^59^. Likewise, uptake of propidium ions across intact cell membranes has been previously observed in bacteria showing high membrane potentials^60^. However, it is important to note that the inherent limitations of this assay include less quantitative assessment of dye penetration and possible toxicity following longer exposure^61^. Our observations suggest that further investigation into the mechanisms underlying transient membrane permeability in different cell types and stress conditions could lead to improved methods for accurately measuring cell viability, ultimately enhancing our understanding of cellular stress.

We report, to our knowledge, the first genome-scale hibernator CRISPR-Cas9 screen in cells derived from Syrian hamster. To create a genome-scale CRISPR-Cas9 library, we implemented a CRISPOR-based computational pipeline for sgRNA selection and benchmarked its success by carrying out the hamster and human screens described here. Our validated sgRNA libraries and associated algorithms should serve as a valuable resource to the hibernation community, as well as other scientific communities developing CRISPR-based tools for non-model organisms.

Our data point to GPX4 levels as a major determinant of cellular cold tolerance. Indeed, we observed that GPX4 protein levels were statistically significantly higher in RPE1 and HeLa cells compared to K562 cells (**Figure S13)**, which correlates with increased cold tolerance. However, beyond this limited analysis, whether endogenous levels of functional GPX4 across various cell types correlates with their cold tolerance remains to be tested. We briefly explored the possibility that GPX4 levels might be regulated by cold exposure and did not observe significant changes in GPX4 mRNA and protein levels in cold-exposed cells (**Figure S14**). It remains possible, however, that GPX4 levels are modulated in animals initiating hibernation. While several studies failed to observe altered *Gpx4* mRNA levels in the context of hibernation^62–65^, at least one study has reported an increase in *Gpx4* transcript levels during hibernation in the heart of a hibernating squirrel^66^. Beyond modulation of GPX4 levels, it also remains possible that other factors such as GPX4 subcellular localization, the abundance of the cellular selenocysteine incorporation machinery, the abundance of glutathione, as well as GPX4-pathway independent factors all play important roles in limiting cold tolerance across various cell types. Although we did not see an increase in cold cell viability upon treatment of K562 cells with GSH-MEE, it is notable that in our screens, sgRNAs targeting genes involved in the glutathione biosynthesis (e.g. *GCLC*, *GSS*^67,68^) showed increased depletion in cold-exposed K562 versus BHK-21 cells (**Figure S12, Figure S15**). This suggests a reduced dependency on the conventional glutathione biosynthetic pathway in cold-exposed BHK-21 cells. In addition, while our data points to a key cellular need to reduce cold-associated lipid hydroperoxides, the mechanisms that drive peroxide accumulation in the cold remain unclear^69,70^. Specifically, the roles of reactive oxygen species (ROS) production, polyunsaturated fatty acid (PUFA) levels, and free iron concentration should be further explored to understand the specific drivers of cold-induced lipid peroxidation and ferroptotic cell death.

Our findings implicating GPX4 in cold tolerance suggest potential therapeutic strategies for enhancing cellular resilience to cold-induced stress. Modulating GPX4 activity or suppressing ferroptosis may be beneficial in contexts such as acute cold injury and organ preservation, where protecting cells from cold-associated oxidative damage is critical^71^. In organ transplantation, for example, tissues are commonly stored at low temperatures to reduce metabolic demand, and approaches that enhance GPX4 activity and/or inhibit ferroptosis could improve tissue viability during preservation.

Our study has several limitations. We focused the analysis of our CRISPR screen on genes that are selectively essential at 4°C versus 37°C, and should note that pathways that affected cellular fitness to a similar extent at 4°C and 37°C were not prioritized. Our screens are designed to knock out a single gene per cell, and thus gene families with redundant functions were likely missed. We limited our current investigations to cell culture paradigms. Although factors such as GPX4, FOXO1, and 5-ALA^24,29^ have been implicated in cold tolerance *in vitro*, future *in vivo* studies in hibernating organisms that naturally undergo profound drops in core body temperature could yield further insight into the adaptive mechanisms that govern cellular survival during periods of hibernation and torpor, while also allowing for the study of cold tolerance mechanisms in cell types that cannot presently be maintained in culture. Similarly, a *Gpx4* conditional knockout mouse line has been generated and used to show that *Gpx4* is essential for mitochondrial integrity and neuronal survival in adult animals^72^; however, these animals have not yet been used to evaluate the contribution of *Gpx4* to *in vivo* cellular and tissue cold tolerance.

The ability to hibernate and survive long-term cold exposure is present in many mammalian species from rodents to bats to primates, raising the possibility that the ability to enter hibernation and tolerate cold was an ancestral trait present in protoendotherms. Our observation that a pathway centered around GPX4 is required across diverse mammalian species to protect cells from cold-induced cell death, raises a question whether GPX4 has a role in protection from cold-induced ferroptosis in other cold-tolerant vertebrates including fish, amphibians, reptiles, and birds that also enter torpor. Such studies will advance our understanding of the evolved mechanisms by which cells mitigate cold-associated damage, one of the most common challenges faced by cells and organisms in nature.

## Acknowledgments

We thank Wei Li for generously sharing 13-lined ground squirrel (*Ictidomys tridecemlineatus*) fibroblasts and Thomas P. Zwaka for sharing greater horseshoe bat (*Rhinolophus ferrumequinum*) fibroblasts. We thank Amanda Chilaka and Sumeet Gupta of the Whitehead Institute Genome Technology Core for high-throughput sequencing. We thank Fabian Schulte and Brooke Linnehan of the Whitehead Institute Quantitative Proteomics Core for mass spectrometry. We thank Aurora Lavin-Peter, Aleksandar Markovski, William S. Owens, Sheamin Khyeam, Karina Smolyar, and Elena Assad for their technical support. We thank Whitney Henry, Matthew Vander Heiden, Alison Ringel, Morgan Sheng, Myriam Heiman, and members of the Hrvatin laboratory for providing feedback on the study. Schematics created with BioRender.com. This material is based upon work supported by the National Science Foundation Graduate Research Fellowship under Grant No. 2141064 and The Paul and Daisy Soros Fellowship for New Americans (PSM). Research reported in this publication was supported by The G. Harold and Leila Y. Mathers Charitable Foundation and the NIDDK/NIH under Award Number DP2DK136123. The content is solely the responsibility of the authors and does not necessarily represent the official views of the National Institutes of Health.

## Author contributions

B.L., K.M.K., and S.H. conceived and designed the study. B.L. and K.M.K. designed, performed, and analyzed experiments. B.L., K.M.K., H.R.K. designed and performed genome wide CRISPR screens. A.K. assisted with the genome wide CRISPR screens. B.L. performed the genetic and pharmacological studies to validate the screens. K.M.K. performed the experiments with primary cells. T.G.T. carried out transcriptomic and proteomic assays, analyzed data, and edited manuscript during revisions. E.M.F. and G.W.B. designed algorithms to select genome-wide Syrian hamster (*Mesocricetus auratus*) CRISPR library. M.E.B. and P.B. immortalized mouse embryonic fibroblasts and greater horseshoe bat (*Rhinolophus ferrumequinum*) fibroblasts. J.M.R., R.J., J.S.W., and E.C.G. advised on the study. B.L., S.H., and E.C.G led the writing of the manuscript with contributions from other authors. E.C.G. and S.H. obtained funding for the research. All authors approved and reviewed the manuscript.

## Tables

**Table S1.** Significant genes from Cycle 3 of genome-wide BHK-21 screen (median log2 fold-change < -1 or > 1 and FDR < 0.1)

| Gene | Median Log2FC | FDR |
| --- | --- | --- |
| Rps29 | 1.3794 | 0.017327 |
| Dld | -1.7915 | 0.003094 |
| Eefsec | -2.1607 | 0.000707 |
| Gpx4 | -2.2215 | 0.000707 |
| Lias | -1.1881 | 0.011251 |
| Lipt1 | -1.1444 | 0.05562 |
| Pstk | -2.3621 | 0.000707 |
| Secisbp2 | -1.1041 | 0.000707 |
| Sepsecs | -2.3116 | 0.000707 |
| Ybey | -1.5789 | 0.017492 |

**Table S3.** Significant genes from 15 days 4°C of genome-wide BHK-21 screen (median log2 fold-change < -0.5 or > 0.5 and FDR < 0.1)

| Gene | Median Log2FC | FDR |
| --- | --- | --- |
| LOC101827392 | 0.5431 | 0.066832 |
| Eefsec | -0.81913 | 0.00165 |
| Gpx4 | -1.5133 | 0.00165 |
| Secisbp2 | -0.74996 | 0.00165 |

**Table S4.**
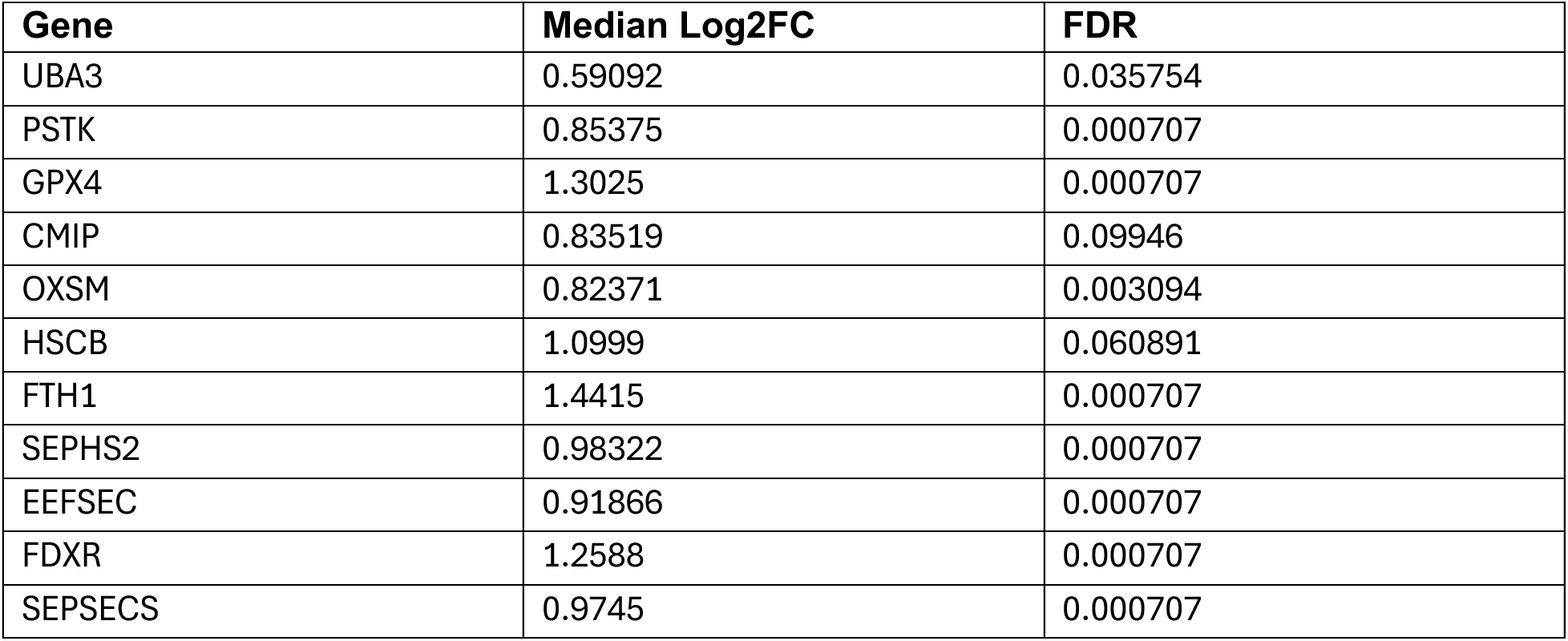
Significant genes from Cycle 3, 4°C + ferrostatin-1 vs 4°C of genome-wide K562 screen (median log2 fold-change < -0.5 or > 0.5 and FDR < 0.1)

| Gene | Median Log2FC | FDR |
| --- | --- | --- |
| UBA3 | 0.59092 | 0.035754 |
| PSTK | 0.85375 | 0.000707 |
| GPX4 | 1.3025 | 0.000707 |
| CMIP | 0.83519 | 0.09946 |
| OXSM | 0.82371 | 0.003094 |
| HSCB | 1.0999 | 0.060891 |
| FTH1 | 1.4415 | 0.000707 |
| SEPHS2 | 0.98322 | 0.000707 |
| EEFSEC | 0.91866 | 0.000707 |
| FDXR | 1.2588 | 0.000707 |
| SEPSECS | 0.9745 | 0.000707 |

## Methods

### Cell culture

Two hibernator-derived (BHK-21, HaK) and four non-hibernator-derived (HT1080, RPE1, HeLa, K562) cell lines were used. All cells were purchased from ATCC. K562 cells were cultured in Roswell Park Memorial Institute (RPMI) 1640 Medium (Gibco #11875093) supplemented with 10% fetal bovine serum (GeminiBio #100-106) and 1% penicillin/streptomycin. The remaining cells were cultured in Dulbecco’s modified Eagle’s medium (DMEM) (Thermo Fisher Scientific #12430054) supplemented with 10% fetal bovine serum (GeminiBio #100-106) and 1% penicillin/streptomycin (Thermo Fisher Scientific #15140122).

### Cell viability assay

Cells were seeded on 24-well culture plates at 50,000 cells per well and allowed to adhere for 24 hours at 37°C. Cells were then placed at 4°C with 5% CO2 for varying time periods (1, 4, or 7 days), with a 30-minute rewarming at 37°C before assessing cell viability via trypan blue (TB) (Thermo Fisher Scientific #15250061) staining. To assess cell viability using the TB assay, cells were washed with PBS (Thermo Fisher Scientific #10010023), trypsinized and centrifuged at 300 x g for 3 minutes. The resulting pellet was resuspended in media, TB was added to a final concentration of 0.2%, and cells were manually counted using a hemocytometer.

### Assessment of Lipid Peroxidation

Cells were seeded on 24-well culture plates at 100,000 cells per well at 37°C. 24h later, cells were then incubated with 5 μM C11 BODIPY (ThermoFisher Scientific #D3861) for 30 minutes at 37°C before washing and media replacement. Cells were then placed at 4°C or 37°C for 1 day. After 1 day, cells were washed, trypsinized, and resuspended in PBS with CountBright™ Absolute Counting Beads (ThermoFisher Scientific #C36950). Samples were analyzed by flow cytometry on the BD LSRFortessa Cell Analyzer.

### Assessment of cell permeability using FACS

Cells were seeded on 24-well culture plates at 100,000 cells per well and cultured for 24h at 37°C. Cells were placed at 4°C or 37°C for 1 day and then incubated with 1 μM TO-PRO-3 (Thermo Fisher Scientific #T3605) for 15 minutes at the incubation temperature (4°C or 37°C) before washing and media replacement. Cells were rewarmed for 30 minutes at 37°C, and stained with 1 μg/mL DAPI (Thermo Fisher Scientific #D1306) for 15 minutes at 37°C. Cells were then washed, resuspended in PBS with CountBright™ Absolute Counting Beads (ThermoFisher Scientific #C36950) and analyzed by flow cytometry on the BD LSRFortessa Cell Analyzer.

### Cell death assay (LDH)

The amount of LDH in the supernatant was measured using the LDH-Glo Cytotoxicity Assay (Promega J2380) following the manufacturer’s protocol. The samples were mixed with reagents on microplates, and luminescence was measured after a 30-minute incubation at room temperature. The maximum amount of LDH in each was measured by fully lysing replicate wells with 2% Triton X-100 (Thermo Fisher Scientific #A16046.AE). Background LDH signal was measured from media-only wells and subtracted from sample values before normalization to fully lysed wells in order to determine the amount of cytotoxicity per sample.

### Compound sources

ML162 (SML0521), RSL3 (SML2234), erastin (E7781), 2’2’-Bipyridyl (D216305), and necrostatin-1 (N9037) were obtained from MilliporeSigma. Ferrostatin-1 (S7243), Liproxstatin-1 (S7699), and z-VAD-FMK (S7023) were purchased from Selleck Chemicals LLC. Deferoxamine mesylate (ab120727) was purchased from Abcam.

### Genome-wide library design

sgRNAs targeting the hamster genome build GCF_017639785.1 (BCM_Maur_2.0) were designed based on NCBI Refseq gene annotations. Protein-coding regions of all gene exons were filtered to retain only the most constitutively expressed exons and extended 20 nt on each end. These regions were provided as input for CRISPOR^73^ to pick and score all potential guides. For each gene, guides were filtered by potential problem flags and then ranked by an overall score derived from its efficiency (the Doench16/Fusi score), a custom motif penalty, a fractional frameshift (from the Lindel score and the fractional distance to the beginning of the region), and its specificity (derived from the MIT specificity score). For each gene, after the guide with the best score was chosen, subsequent guides were also penalized by a location score, adding a penalty for presence in the same exon and proximity to previously selected guides for the same gene. If more than ten potential guides were identified for a gene, the ten with the highest scores were selected.

For the human genome-wide library, we obtained a list of protein-coding genes in human based on Ensembl release 98. The top five sgRNAs were picked for protein-coding genes and controls. Human intergenic sgRNAs were chosen by first subsetting the list of all designed intergenic-region-targeted sgRNAs by Rank = 1, then requiring the sgRNAs to target only one genomic locus. This list of 465 sgRNAs was ranked by MIT specificity score in descending order, and the top 449 sgRNAs were chosen. The final library contained a total of 98,077 unique sgRNAs.

### Library cloning

For the hamster library, 214,116 unique protospacer sequences targeting ∼21,000 unique gene symbols (> 99% with 10 sgRNAs/gene symbol), along with 2,299 intergenic-targeting and 250 nontargeting sequences were synthesized as an oligonucleotide pool (Agilent Technologies).

Separately, for the human library, 98,077 unique protospacer sequences targeting ∼19,600 Ensembl transcript IDs (> 99.9% with 5 sgRNAs/gene), 449 intergenic-targeting sequences, 50 non-targeting sequences, and one sequence targeting the *AAVS1* safe harbor locus were synthesized as an oligonucleotide pool (Agilent Technologies). A guanine nucleotide was prepended to each 20-nucleotide protospacer sequence that began with A, C, or T. For both hamster and human libraries, homology arms were prepended (5’-TATCTTGTGGAAAGGACGAAACACC-3’) and appended (5’-GTTTAAGAGCTATGCTGGAAACAGCATAGC-3’). Additional adapters were pre- and appended to the hamster library to enable subpool amplification if desired (subpool 1: 5’-TCGGTGTATTGCTAGTGCGAACCCA-3’ and 5’-ATCGTGTGAAAGGTGCCGCTATTGC-3’; subpool 2: 5’-GGTTCGTTCTACACATGGAAGCGGC-3’ and 5’-GAGGACTTCGAGTAGAACGCTGGCG-3’). Oligonucleotide pools were PCR amplified (forward primer: 5’-TTTCTTGGCTTTATATATCTTGTGGAAAGGACGAAACACC-3’; reverse primer: 5’-ATTTAAACTTGCTATGCTGTTTCCAGCATAGCTCTTAAAC-3’) and cloned into pLentiCRISPRv2-Opti (a gift from David Sabatini; Addgene plasmid # 163126; http://n2t.net/addgene:163126; RRID:Addgene_163126). Briefly, 0.25 µL of a 100 nM pool was PCR amplified in 50 µL reactions using Q5 HotStart DNA Polymerase (New England Biolabs) for 10 cycles under the following conditions:

|  |  |  |
| --- | --- | --- |
| 1 cycle | 98°C | 2 minutes |
| 10 cycles | 98°C | 10 seconds |
|  | 55-62°C | 15 seconds |
|  | 72°C | 15 seconds |
| 1 cycle | 72°C | 2 minutes |
| 1 cycle | 4°C | hold |

All concentration steps were performed using a DNA Clean and Concentrator-5 kit (ZymoResearch). Amplification from 8 gradient annealing reactions (55-62°C) was assessed, and successful reactions were pooled and concentrated. The vector was digested overnight at 37°C with FastDigest Esp3I and FastAP (ThermoFisher Scientific), gel purified using a Zymoclean gel DNA recovery kit (ZymoResearch), and concentrated. NEBuilder HiFi DNA Assembly Master Mix (New England Biolabs) was used in 2 x 40-80 µL bulk assembly reactions, for a combined total reaction volume of 160 µL containing 4 µg of vector and 160 ng of PCR amplicon (hamster) or 120 µL total containing 3 µg of vector and 120 ng of PCR amplicon (human). Each bulk reaction was distributed in 5 µL aliquots and incubated for 10 minutes at 52.2°C, pooled, concentrated, introduced into Endura Electrocompetent DUO cells (Lucigen) by electroporation, and plated on 8-16 LB agar with 75 µg/mL carbenicillin in 245 mm x 245 mm square bioassay dishes. A dilution series was also plated to assess electroporation efficiency. Cells were incubated overnight at 30°C, collected, and DNA was isolated using a ZymoPURE II Plasmid DNA Maxiprep kit (ZymoResearch). Plasmid from separate electroporations was combined proportionally based on electroporation efficiency for a combined total library coverage of > 50-fold for each library. Sequence representation in the libraries was assessed as described below.

### Lentivirus production for CRISPR-Cas9 screen

For large-scale virus production, HEK-293T (1.5×10^7^) cells were seeded in T175 cm^2^ flasks in DMEM (Thermo Fisher Scientific #12430054) supplemented with 10% fetal bovine serum (GeminiBio #100-106). Media was changed 24 hours later to 20 mL viral production medium: IMDM (Thermo Fisher Scientific #1244053) supplemented with 20% heat-inactivated fetal bovine serum (GeminiBio #100-106). Cells were transfected 8 hours later with a mix containing 76.8 µL Xtremegene-9 transfection reagent (MilliporeSigma #06365779001), 3.62 µg pCMV-VSV-G (Addgene plasmid # 8454; http://n2t.net/addgene:8454; RRID:Addgene_8454)^74^, 8.28 µg psPAX2 (a gift from Didier Trono; Addgene plasmid # 12260; http://n2t.net/addgene:12260; RRID:Addgene_12260), and 20 µg sgRNA/Cas9 plasmid and Opti-MEM (Thermo Fisher Scientific #11058021) to a final volume of 1 mL. Media was changed 16 hours later to 55 mL fresh viral production medium. Viral supernatant was collected and filtered through a 0.45 µm filter and aliquoted 48 hours post-transfection, then stored at -80°C until use.

### CRISPR screen in K562 cells

K562 cells (3.9×10^8^) were transduced with a pooled genome-wide lentiviral sgRNA library in a Cas9-containing vector at MOI <1. The transduced cells were selected with 3 µg/mL puromycin (Thermo Fisher Scientific #A1113803), and 1×10^8^ cells were passaged every 48-72 hours at a density of 6×10^5^ cells/T175 cm^2^ flask in 45 mL RPMI 1640 (Gibco #11875093) medium supplemented with 10% fetal bovine serum (GeminiBio #100-106) and 1% penicillin/streptomycin for the duration of the screen. At 5 days post-puromycin selection, 1×10^8^ cells were exposed to the cold with or without 1 µM ferrostatin-1 (Selleck Chemicals #S7243). K562 cells (1×10^8^) were collected from the each of surviving populations after cold exposure and/or rewarming as well as a matched untreated population. Genomic DNA was isolated using the QIAmp DNA Blood Maxiprep kit (Qiagen # 51192), and high-throughput sequencing libraries were prepared.

### CRISPR screen in BHK-21 cells

BHK-21 cells (8.55×10^8^) were transduced with a pooled genome-wide lentiviral sgRNA library in a Cas9-containing vector at an MOI of 0.25. Transduced cells were selected with 2 µg/mL puromycin (Thermo Fisher Scientific #A1113803), and 2.56×10^8^ cells were passaged every 48-72 hours at a density of 5×10^6^ cells/15-cm dish in DMEM (Thermo Fisher Scientific #12430054) medium supplemented with 10% fetal bovine serum (GeminiBio #100-106) and 1% penicillin/streptomycin for the duration of the screen. At 6 days post-puromycin selection, 2.56×10^8^ cells were exposed to the cold. Cells (2.4×10^8^) were collected from each of the surviving populations after cold exposure and/or rewarming as well as from a matched untreated population. During the cycles, cells were placed at 4°C for 4 days, then rewarmed to 37°C for 24 hours, reseeded, and after a further 24 hours at 37°C were placed back at 4°C to begin a subsequent cycle. Genomic DNA was isolated using the QIAmp DNA Blood Maxiprep kit (Qiagen # 51192), and high-throughput sequencing libraries were prepared. Cells from long-term cold exposure arms (non-cycling) were also rewarmed for ∼30 minutes prior to being harvested. As such, all cells collected from each arm were those that were attached on the plate. Any cells that did not adhere after 24 hours or 30 minutes (depending on the screen arm paradigm) were discarded prior to collection. To harvest cells, the media was removed and cells were washed with 1XPBS prior to mild (3-5 minute) 0.1% trypsin treatment and subsequent quenching with media and collection by centrifugation.

### Sequencing library preparation

The QIAamp DNA Blood Maxiprep Kit (Qiagen # 51192) was used to extract genomic DNA (gDNA) from cell pellets of 2.5-5×10^7^ cells according to manufacturer’s instructions with minor modifications: QIAGEN Protease was replaced with 500 µL of 10 mg/mL Proteinase K (MilliporeSigma # 3115879001) in water, and cells were lysed overnight; centrifugation was performed for 2 minutes and 5 minutes after the first and second wash, respectively; 1 mL of water preheated to 70°C was used to elute gDNA, followed by centrifugation for 5 minutes. gDNA was quantified using the Qubit dsDNA HS Assay kit (Thermo Fisher Scientific #Q32851). PCR amplification of sgRNA sequences was performed in 50 µL reactions using ExTaq Polymerase (Takara Bio #RR001B) with the following program:

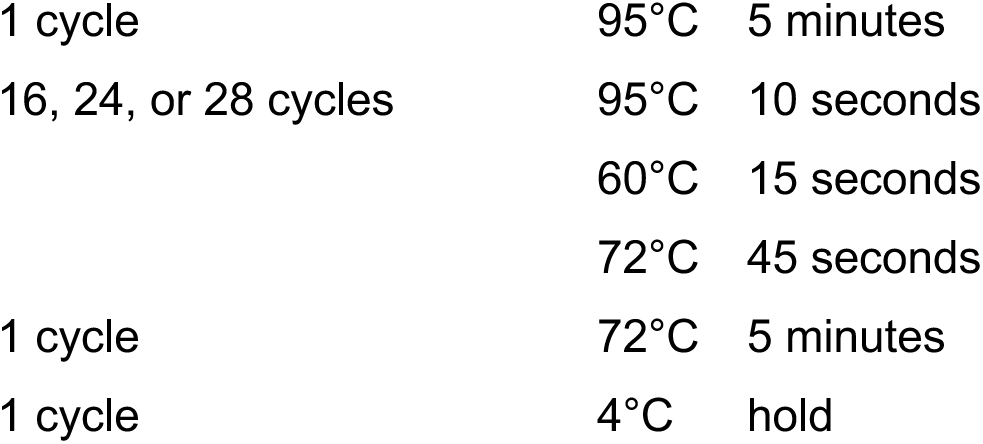

|  |  |  |
| --- | --- | --- |
| 1 cycle | $95^\circ\text{C}$ | 5 minutes |
| 16, 24, or 28 cycles | $95^\circ\text{C}$ | 10 seconds |
| | $60^\circ\text{C}$ | 15 seconds |
| | $72^\circ\text{C}$ | 45 seconds |
| 1 cycle | $72^\circ\text{C}$ | 5 minutes |
| 1 cycle | $4^\circ\text{C}$ | hold |

Using the following primers:

Forward: 5’-AATGATACGGCGACCACCGAGATCTACACCCCACTGACGGGCACCGGA - 3’

Reverse: 5’-CAAGCAGAAGACGGCATACGAGATCnnnnnnTTTCTTGGGTAGTTTGCAGTTTT - 3’

Where “nnnnnn” denotes the barcode used for demultiplexing.

Plasmid libraries were amplified for 16 cycles using 10 ng input per 50 µL reaction. gDNA was initially amplified for 24 (human) or 28 (hamster) cycles in 50 µL test PCR reactions with 1, 3, 4, 5, or 6 µg input. An additional 20-75 reactions were performed using 3-6 µg per reaction (sampling 150 µg or 300 µg gDNA total for human and hamster screens, respectively). Select-a-Size DNA Clean and Concentrator (Zymo Research #D4080) or HighPrep PCR (MagBio Genomics) was used to purify 95-100 µL of each pooled sample, which were eluted with 12-20 µL water and quantified using the Qubit dsDNA HS Assay kit prior to sequencing for 50 cycles on an Illumina Hiseq 2500 or NovaSeq using the following primers:

Read 1 sequencing primer: 5’-GTTGATAACGGACTAGCCTTATTTAAACTTGCTATGCTGTTTCCAGCATAGCTCTTAAAC - 3’Index sequencing primer: 5’- TTTCAAGTTACGGTAAGCATATGATAGTCCATTTTAAAACATAATTTTAAAACTGCAAACTACCCAAGAAA - 3’

### CRISPR screen data analysis

Sequencing reads from the human screen were trimmed and mapped to the sgRNA library using Bowtie version 1.0.0^75^, allowing no mismatches, and counted. Sequencing reads from the hamster screen were trimmed and mapped to the sgRNA library using a combination of command line prompts and a custom Python function which generated a dataframe of counts which perfectly (no permissibility for any base mismatches) matched sequenced reads to the full guide library). Data from both human and hamster screens was similarly analyzed using MAGeCK version 0.5.9.5^76^ using a gene test false discovery rate (FDR) threshold of 0.05, the FDR method for p-value adjustment, and the median as the gene-level scoring metric.

PANTHER overrepresentation test was conducted on a set of 204 genes (FDR < 0.1) from the third cycle of the human K562 CRISPR-Cas9 screen [‘Analyzed List’]. A list of genes expressed in K562 cells at 37°C collected from bulk RNA sequencing was used as the [‘Reference List’]: https://pantherdb.org/tools/compareToRefList.jsp. Fisher’s Exact test type and false discovery rate correction were utilized. Bulk RNA sequencing of K562 cells at 37°C was conducted by generating a cDNA library prepared by depleting rRNA using the NEBNext rRNA Depletion Kit (Human/Mouse/Rat) (New England Biolabs) and then the NEBNext Ultra II Directional RNA Library Prep Kit (New England Biolabs). cDNA libraries were amplified using appropriate multiplexed index primers for PCR.

Data was visualized using R version 4.2.1^77^ corrplot package version 0.92^78^ and base graphics, and GraphPad Prism version 10.

### Generation of stable CRISPR/Cas9-targeted cell lines

Homology arms were prepended (5’-TATCTTGTGGAAAGGACGAAACACC-3’) and appended (5’-GTTTAAGAGCTATGCTGGAAACAGCATAGC-3’) to the top two ranked human or hamster *GPX4*-targeting guides from the genome-wide screens. These guides were PCR amplified and cloned into pLentiCRISPRv2-Opti (a gift from David Sabatini; Addgene plasmid # 163126; http://n2t.net/addgene:163126; RRID:Addgene_163126). K562 and BHK-21 cells were then infected with LentiCRISPRv2. Transduced cells were selected with 3 µg/mL puromycin (Thermo Fisher Scientific #A1113803) for K562 cells and 2 µg/mL for BHK-21 cells beginning two days after infection. After one passage, cells were sorted (BD FACSAria™ Fusion Flow Cytometer) into 96-well plates, at one cell per well to generate clonal knockout cell lines. Cells were cultured in media with 2.5 µM liproxstatin-1 (Selleck Chemicals #S7699) to prevent cell death. Western blotting was used to confirm GPX4 loss in individual knockout clones prior to further use.

Human sgRNA:

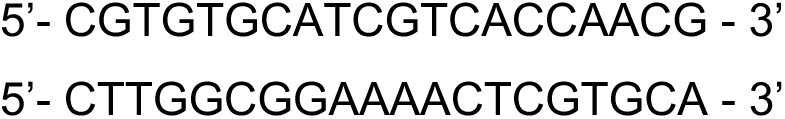

Hamster sgRNA:

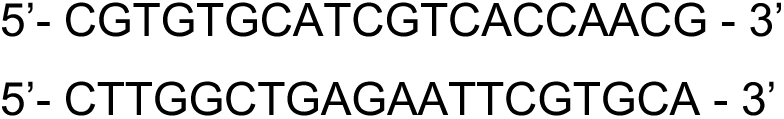

### Generation of CRISPR/Cas9-targeted human kidney fibroblast pooled cells

Primary human kidney fibroblasts were infected with LentiCRISPRv2 with sgRNAs targeting *GPX4*, using the top two ranked *GPX4*-targeting guides from the genome-wide human K562 screen. Transduced cells were selected with 1 µg/mL puromycin (Thermo Fisher Scientific #A1113803) beginning four days after infection. Cells were cultured in media with 2.5 µM liproxstatin-1 (Selleck Chemicals #S7699) to prevent cell death. Western blotting was used to confirm GPX4 loss in individual knockout clones prior to further use.

### Immunoblotting

Cells were plated at 6×10^5^ cells per well in a 6-well plate and maintained overnight at 37°C. Cells were then collected when wells were confluent. The media was aspirated, and cells were washed with ice-cold PBS (Thermo Fisher Scientific #10010023). Cells were scraped in 100 µL of RIPA buffer (150 mM NaCl, 1% Triton X-100, 0.5% Na-Dexycholate, 0.1% SDS, 50 mM Tris, pH 8) with protease inhibitors (MilliporeSigma #04693159001) and collected in microcentrifuge tubes. The contents were then vortexed for 30 minutes at 4°C and centrifuged at 16000 x g for 20 minutes at 4°C. Protein content of the samples was measured using the Pierce BCA Protein Assay kit (Thermo Fisher Scientific #23225) and resuspended in 100 µL of RIPA buffer with protein loading dye (10% SDS, 500 mM DTT, 50% Glycerol, 250 mM Tris-HCl and 0.5% bromophenol blue dye, pH 6.8). Protein samples were boiled for 5 minutes at 95°C, vortexed, and centrifuged at 16000 x g for 5 minutes before loading onto a gel. The gel was run at 120 V for 2 hours before being transferred to a Polyvinylidene difluoride (PVDF) membrane (Thermo Fisher Scientific #88518). The transfer cassette was run at 45 V for 2 hours. The membranes were then transferred to 5% milk TBST (1X Tris-Buffered Saline, 0.1% Tween® 20 Detergent) and shaken for 15 minutes before being washed in TBST. Primary antibodies recognizing GPX4 (Abcam #ab41787) or HA (Cell Signaling Technology #3724) were added at 1:1000 in 5% BSA (MilliporeSigma #A9418) and incubated overnight at 4°C. The membrane was then washed 3 times in 1x TBST and 1:3000 anti-rabbit secondary antibody (Cell Signaling Technology #7074) in 5% milk TBST was added. Membranes were incubated for 1 hour at room temperature and washed 3x in 1x TBST. Membranes were then incubated in Pierce™ ECL Western Blotting Substrate (Thermo Fisher Scientific #32106) for 5 minutes before imaging.

### Generation of HA-GPX4 and HA-GFP overexpression constructs

To generate GPX4-overexpression lines, entire human *GPX4* (cytosolic; NM_001367832.1) and hamster *Gpx4* were amplified from the cDNA of K562 and BHK-21 cells, respectively. The cloned transcript fragments include the 3’ UTR that contains the SECIS element necessary for selenocysteine incorporation. The *GPX4* genomic DNA fragments were cloned into pLJM1 (Addgene plasmid # 19319; http://n2t.net/addgene:19319; RRID:Addgene_19319) and confirmed via sequencing. To create GPX4 mutants, (U46S), non-overlapping primers were used to create mutations via inverse PCR, as previously described^79^, converting the UGA stop codon to UCA, encoding serine. GFP was amplified from pLJM1-EGFP (Addgene plasmid # 19319; http://n2t.net/addgene:19319; RRID:Addgene_19319) with Gibson overhangs to enable insertion into a lentivirus vector. HA-GPX4 and HA-GFP DNA fragments were then amplified by PCR and cloned into blasticidin lentiviral vectors [gift from Whitney Henry at the Whitehead Institute for Biomedical Research] using standard cloning methods, with point mutations in the PAM and guide sequences to prevent targeting of exogenously expressed *GPX4*.

HEK-293T cells (5×10^5^) were seeded in 2 wells of a 6-well plate in DMEM (Thermo Fisher Scientific #12430054) supplemented with 10% fetal bovine serum (GeminiBio #100-106). Media was changed 16 hours later to 2 mL per well of viral production medium: IMDM (Thermo Fisher Scientific #1244053) supplemented with 20% heat-inactivated fetal bovine serum (GeminiBio #100-106). Cells were transfected 8 hours later with a mix containing 4.2 µL Xtremegene-9 transfection reagent (MilliporeSigma #06365779001), 181 ng pCMV-VSV-G (Addgene plasmid # 8454; http://n2t.net/addgene:8454; RRID:Addgene_8454)^74^, 414 ng psPAX2 (a gift from Didier Trono; Addgene plasmid # 12260; http://n2t.net/addgene:12260; RRID:Addgene_12260), and 1 µg HA-GPX4 or HA-GFP plasmid and Opti-MEM (Thermo Fisher Scientific #11058021) to a final volume of 50 µL per well. Media was changed 16 hours later to 2.5 mL per well of fresh viral production medium. Viral supernatant was collected and aliquoted 48 hours post-transfection, then stored at -80°C until use.

K562 or BHK-21 cells (6×10^5^, wild-type and GPX4 knockout clones) were transduced with the lentiviral HA-tagged GPX4 or GFP in the presence of 10 µg/mL polybrene. After 8 hours of incubation at 37°C, media was changed to virus-free media. After 48 hours, the transduced cells were selected with blasticidin (Thermo Fisher Scientific #A1113903) at a concentration of 6 µg/mL for K562 cells and 4 µg/mL for BHK-21 cells. Overexpression of GPX4 and GFP was confirmed via immunoblotting.

### Primary cell culture

Three hibernator (13-lined ground squirrel postnatal dermal fibroblasts [gift from Wei Li at Duke University], SV40-immortalized Greater horseshoe bat embryonic dermal fibroblasts [gift from Rudolf Jaenisch at the Whitehead Institute for Biomedical Research], and Syrian golden hamster embryonic primary dermal fibroblasts [isolated]) and five non-hibernator (mouse embryonic SV40-immortalized dermal fibroblasts [gift from Jonathan Weissman at the Whitehead Institute for Biomedical Research], adult human kidney fibroblasts [purchased from ATCC], adult human dermal fibroblasts [purchased from ATCC], adult rat dermal fibroblasts [purchased from ATCC], and adult mouse fibroblasts [purchased from ATCC]) primary cells were cultured in Dulbecco’s modified Eagle’s medium (Thermo Fisher Scientific #12430054) supplemented with 10% fetal bovine serum (GeminiBio #100-106) and 1% penicillin/streptomycin (Thermo Fisher Scientific #15140122). Cells were seeded on 24-well culture plates at ∼50,000 cells per well and allowed to adhere for 24 hours at 37°C. Cells were then placed at 4°C with 5% CO_2_ for varying time periods (1, 4, or 7 days), with a 30-minute rewarming at 37°C before assessing cell viability via trypan blue (TB) (Thermo Fisher Scientific #15250061) staining. To assess cell viability using the TB assay, cells were washed with PBS (Thermo Fisher Scientific #10010023), trypsinized and centrifuged at 300 x g for 3 minutes. The resulting pellet was resuspended in media, TB was added to a final concentration of 0.2%, and cells were manually counted using a hemocytometer.

### Primary neuron isolation and culture

Primary Syrian golden hamster and C57BL/6 mouse cortical neurons were isolated at postnatal day 0. The day before isolation, standard tissue culture treated plates were coated with 3 µg/mL Poly-L-Ornithine (MilliporeSigma #P4957) and warmed at 37°C for 24 hours. The morning of isolation, wells were washed with 1X PBS (Thermo Fisher Scientific #10010023). To isolate neurons, pups were placed on ice for 2-5 minutes until movement ceased. Afterwards, they were washed with 10% ethanol, decapitated, and their heads placed in a 10x dissociation media bath (10.16 g MgCl_2_ hexahydrate and 11.915 g HEPES in 450 mL HBSS brought to a pH of 7 using NaOH with subsequent addition of 1.182 g of kynurenic acid heated to 65°C while stirring until the solution was fully dissolved and cooled to room temperature and adjusted to a pH of 7.2. The dissociation media was then diluted from 10X to 1X in HBSS and filtered through a 0.22-µm vacuum filter). Dissected brains were placed in fresh dissociation media, where the cortices were dissected out and transferred to 50 mL Falcon tubes on ice containing 1X dissociation media. Afterwards, the dissociation media was removed from the cortices and replaced with papain solution (5 mL 1x dissociation media, 5-7 grains of L-Cysteine, and 172 µL Worthington papain [crystalline suspension in 50mM sodium acetate, pH 4.5 incubated in 1.1mM EDTA, 0.067mM mercaptoethanol and 5.5mM cysteine-HCl at 37°C. Activates to 20 units per 1 mg protein]) that was preheated to 37°C for 30 minutes. The cortices were incubated in the papain solution for 3-5 minutes at 37°C. Afterwards, the papain solution was replaced with trypsin inhibitor solution (10 mg of trypsin inhibitor dissolved in 10 mL of 1X dissociation media, preheated to 37°C) and incubated subsequently at 37°C for 3-5 minutes, repeated 3 times.

Afterwards, the trypsin inhibitor solution was carefully removed and replaced with 6 mL of preheated complete neurobasal media (5 mL of GlutaMAX5, 10 mL of B-27, 5 mL of penicillin/streptomycin in neurobasal media filtered through a 0.22-µm vacuum filter). Using a 5 mL pipette tip, cortices were triturated until no tissue clumps were present. The cortices were then passed through a 100-µm cell strainer to remove large debris and centrifuged at 300 x g for 3 minutes before resuspension in 5 mL of complete neurobasal media before plating at a density of 50,000 mixed cortical neuron isolated cells per well of a 24 well plate. Subsequently, 50% of the media was replaced every second day with fresh 37°C pre-warmed media.

### RNA-seq

Cell lines were maintained at 37°C, 5% CO₂, and 95% relative humidity. For cold exposure experiments, cells were subsequently transferred to 4°C for up to 5 days. At the time of collection, cells were collected (with gentle scraping for adherent BHK-21 lines) and pelleted by centrifugation. Cell pellets were lysed and homogenized using QIAshredder columns (Qiagen), and RNA was extracted using the RNeasy Mini Kit (Qiagen) with on-column DNase digestion, according to the manufacturer’s instructions. Libraries were prepared using the NEBNext Ultra II kit. In brief, 1 μg of RNA per sample underwent Poly(A) selection before fragmentation and random priming followed by first- and second-strand complementary DNA synthesis. Following cDNA synthesis, NEBNext Multiplex Oligos for Illumina (96 Unique Dual Index Primer Pairs, NEB E6441A) were ligated before PCR enrichment. All libraries were pooled to a final concentration of 2 nM and sequenced on a NovaSeq 6000.

### Proteomics Sample preparation

BHK-21 hamster fibroblast cells were cultured at either standard 37°C conditions or 4°C for 4 days (n = 4 biological replicates per condition, 8 samples total). Cells were lysed at 70°C for 15 min in 1X SDS lysis buffer (5% (w/v) SDS, 100 mM TEAB pH 8.5, 40 mM CAA, 10 mM TCEP, protease inhibitor cocktail, phosphatase inhibitor cocktail). Cellular debris was removed by centrifugation at 15,000 × g for 5 min. Protein (600 µg) was purified via the SP3 method (Hughes et al.). Proteins were bound to 120 µL SP3 beads by adding 2.4 mL 100% (v/v) ethanol, followed by three washes with 1 mL 80% (v/v) ethanol. Samples were digested with trypsin/LysC mix (1:50) in 300 µL 100 mM TEAB overnight in a shaking incubator at 37°C, 115 RPM. The following day, an additional 3 µg trypsin/Lys-C mix in 3 µL 100 mM TEAB was added and digestion continued for 4 hours at 37°C. Peptide digests (300 µg) were dried in a speed-vac concentrator and resuspended in 70 µL 200 mM TEAB.

### TMT labeling and fractionation

Peptides were labeled with TMT10-plex isobaric tags (8 of the 10 available reporter channels used: 126, 127N, 127C, 128N, 128C, 129N, 129C, 130N — 4× 37°C, 4× 4°C). Labeled samples were quenched for 30 min at room temperature with 5% hydroxylamine in 100 mM TEAB. Labeled peptides were pooled together and desalted on a C18 SepPak column. The pooled, labeled peptide sample was fractionated using a Strong Anion Exchange (SAX) column, with increasing concentrations of ammonium acetate (0, 20, 50, 100, 200, 500 mM) used for elution. Fractions were pooled into two sets based on salt concentration:

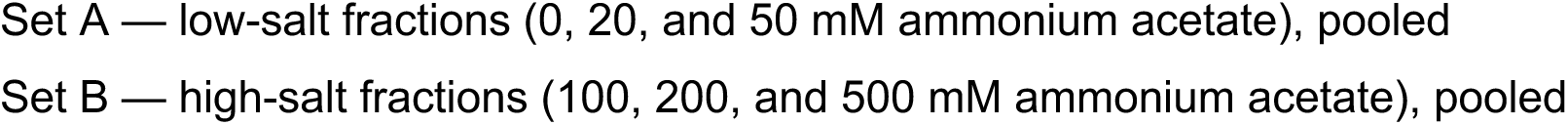

Each pool (Set A and Set B) was lyophilized, then separately fractionated using a High pH Reversed-Phase Peptide Fractionation Kit with a 12-step gradient of increasing acetonitrile concentration (5, 7.5, 10, 12.5, 15, 17.5, 20, 22.5, 25, 27.5, 30, 60%). The resulting 12 fractions from each pool were concatenated pairwise into 6 fractions each (1+7, 2+8, 3+9, 4+10, 5+11, 6+12) and lyophilized, yielding 12 total fractions (6 from Set A, 6 from Set B) for LC-MS/MS. Lyophilized fractions were resuspended in 12 µL 0.2% formic acid in MS-grade water.

### LC-MS/MS data collection

Mass spectrometry was performed using an Orbitrap Eclipse mass spectrometer equipped with a FAIMS Pro source, connected to a Vanquish Neo nLC chromatography system using an EasySpray ES902 column (75 µm × 25 cm, 100 Å) (all Thermo Fisher Scientific, Waltham, MA, USA). Peptides were separated at 300 nL/min on a gradient of 3-25% B for 90 min, 25-40% B for 30 min, 40-95% B for 10 min, and 95% B for 6 min (A: 0.1% FA in water; B: 0.1% FA in 80% acetonitrile). The Orbitrap and FAIMS Pro were operated in positive ion mode (2100 V ion voltage, 305°C ion transfer tube temperature, 4.2 L/min carrier gas flow; FAIMS compensation voltages -45, -55, -65 V). Full scan spectra were acquired in profile mode at 120,000 resolution (MS1 and MS2), 400-1400 m/z scan range, 200 ms maximum fill time, AGC target 100% (MS1) / 250% (MS2), 0.7 m/z isolation window, 2.0e4 intensity threshold, 2-6 charge state, 60 s dynamic exclusion, and 35% HCD collision energy.

### Proteomics database search and protein identification

Raw MS data were processed with PEAKS Studio (version 10.6). Spectral preprocessing included automatic centroiding, deisotoping, and deconvolution. Peptides and proteins were identified by searching against the *Mesocricetus auratus* (Syrian hamster) reference proteome [Uniprot-ID: 10036]. The precursor (parent ion) mass tolerance was set to 15 ppm and the fragment ion mass tolerance to 0.05 Da. Trypsin was specified as the protease with specific cleavage, excluding cleavage at K/R–P sites, and up to three missed cleavages were allowed. Methionine oxidation and serine/threonine/tyrosine phosphorylation were set as variable modifications, and cysteine carbamidomethylation and TMT labeling as fixed modifications, with a maximum of three variable modifications per peptide. The false discovery rate (FDR) was estimated using a decoy-fusion approach, and identifications were filtered to a peptide score of –10lgP ≥ 20 and a 1% FDR at both the peptide and protein levels. Quantified results were exported as .csv files and processed further as described previously^80^.

### Statistical analyses and software information

Data are generally plotted as mean ± S.E.M. unless otherwise indicated. No statistical methods were used to predetermine sample sizes. Unless otherwise indicated, all replication numbers in the figure legends (n) indicate biological replicates. Statistical significance was determined using a two-tailed student’s t-test, one-way ANOVA, or two-way ANOVA using Prism 10 software (GraphPad Software) unless otherwise indicated. Statistical significance was set at p<=0.05 unless otherwise indicated. Figures were finalized in Adobe Illustrator 2024.

## Data availability

Raw and processed RNA-sequencing data generated in this study have been deposited in the NCBI Gene Expression Omnibus (GEO) under accession GSE339558. Mass-spectrometry proteomics data have been deposited to the ProteomeXchange Consortium via the PRIDE partner repository with the dataset identifier PXD081500.

## Supplemental information

Document S1. Figures S1-S15

Table S2. Excel file containing hamster screen data, related to Figure 2

Table S5. Excel file containing human screen data, related to Figure 4

Table S6. Excel file containing GO_Biological Processes, related to Figure S8

## Supplement

**Supplement 1.**
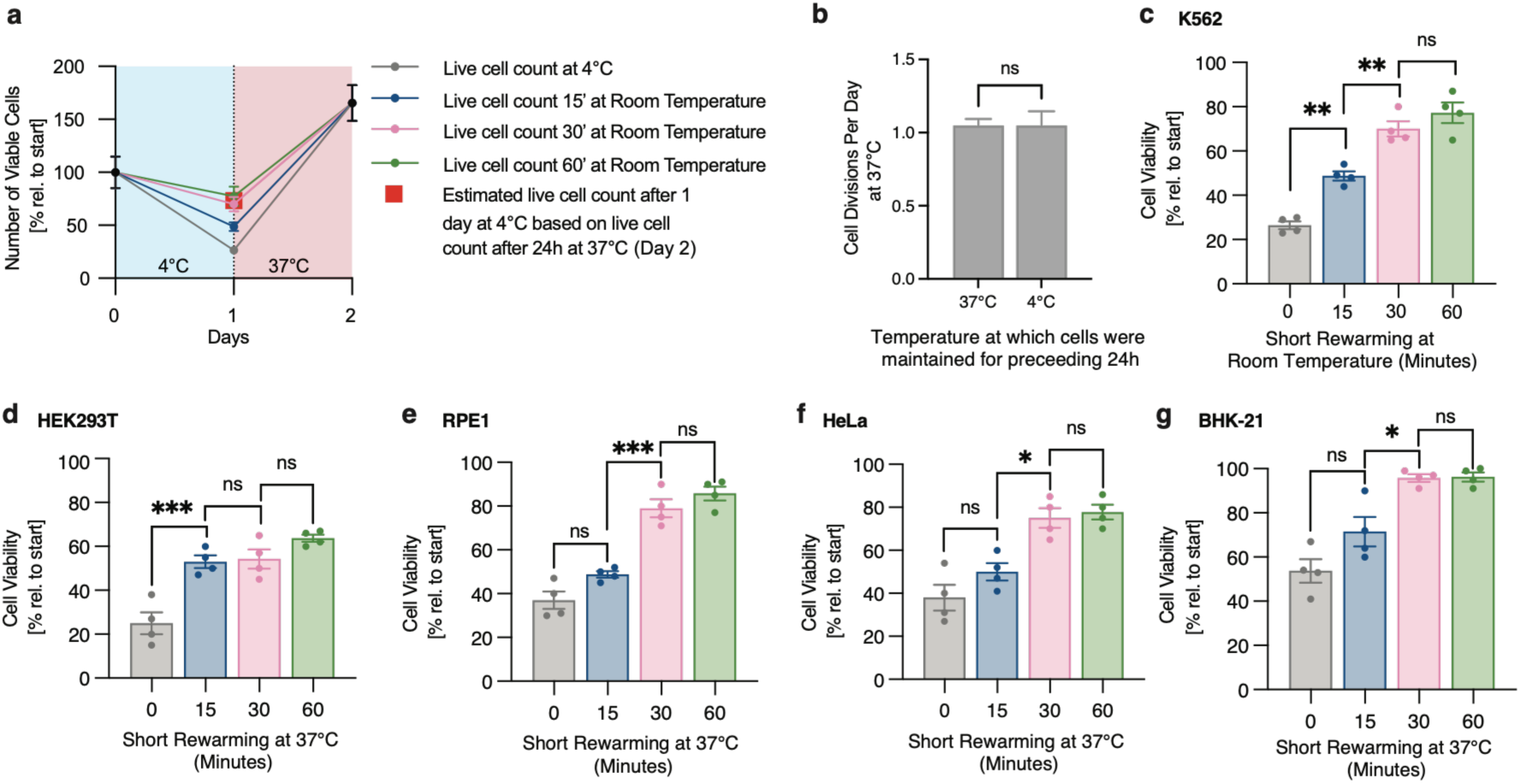
Permeability to trypan blue changes rapidly upon cell rewarming. **a**, Number of viable K562 cells based on trypan blue staining after one day at 4°C and subsequent rewarming for 24 hours at 37°C. Numbers are normalized to initial cell counts. Dots indicate viable cell number based on trypan blue staining of cells after incubation at room temperature for 15, 30, or 60 minutes. Blue shaded regions indicate 4°C exposure and shaded red regions indicate 37°C exposure. Red square indicates calculated cell counts after one day at 4°C based on the viable cell number measured after 24-hour rewarming. **b,** Cell division rates at 37°C of cells continuously maintained at 37°C (left column, n = 3) and of cells exposed to 4°C for 24 h and then transferred to 37°C for 24 h (right column, n = 4). Values show mean ± SEM, with significance measured by student’s t-test. ns P>0.05. **c,** Viability of K562 cells was assessed by trypan blue staining after incubation at room temperature for 0, 15, 30, or 60 minutes following 24 hours at 4°C (*n* = 4). Cells incubated at room temperature for 15 minutes show a significant increase in cell counts compared to cells counted immediately (\*\**P* = 0.0016). Cells incubated at room temperature for 30 minutes show a significant increase in cell counts compared to a 15-minute incubation (\*\**P* = 0.0023), while no significant difference in viability was observed between cells incubated for 30 or 60 minutes (ns, *P* = 0.4059). **d-g**, Viability of cells was assessed by trypan blue staining after incubation at 37°C for 0, 15, 30, or 60 minutes following 24 hours at 4°C (*n* = 4). **d**, HEK293T, **e**, RPE1, **f**, HeLa, **g**, BHK-21. No significant difference in cell viability was observed between cells incubated for 30 or 60 minutes for HEK293T (ns, *P* = 0.3122), RPE1 (ns, *P* = 0.5137), HeLa (ns, *P* = 0.9735), and BHK-21 (ns, *P* = 0.9998) cells. All values show mean ± SEM, with significance determined by one-way ANOVA adjusted for multiple comparisons by Tukey’s HSD. \**P* < 0.05; \*\**P* < 0.01; \*\*\**P* < 0.001; \*\*\*\**P* < 0.0001; ns *P* > 0.05.

**Supplement 2.**
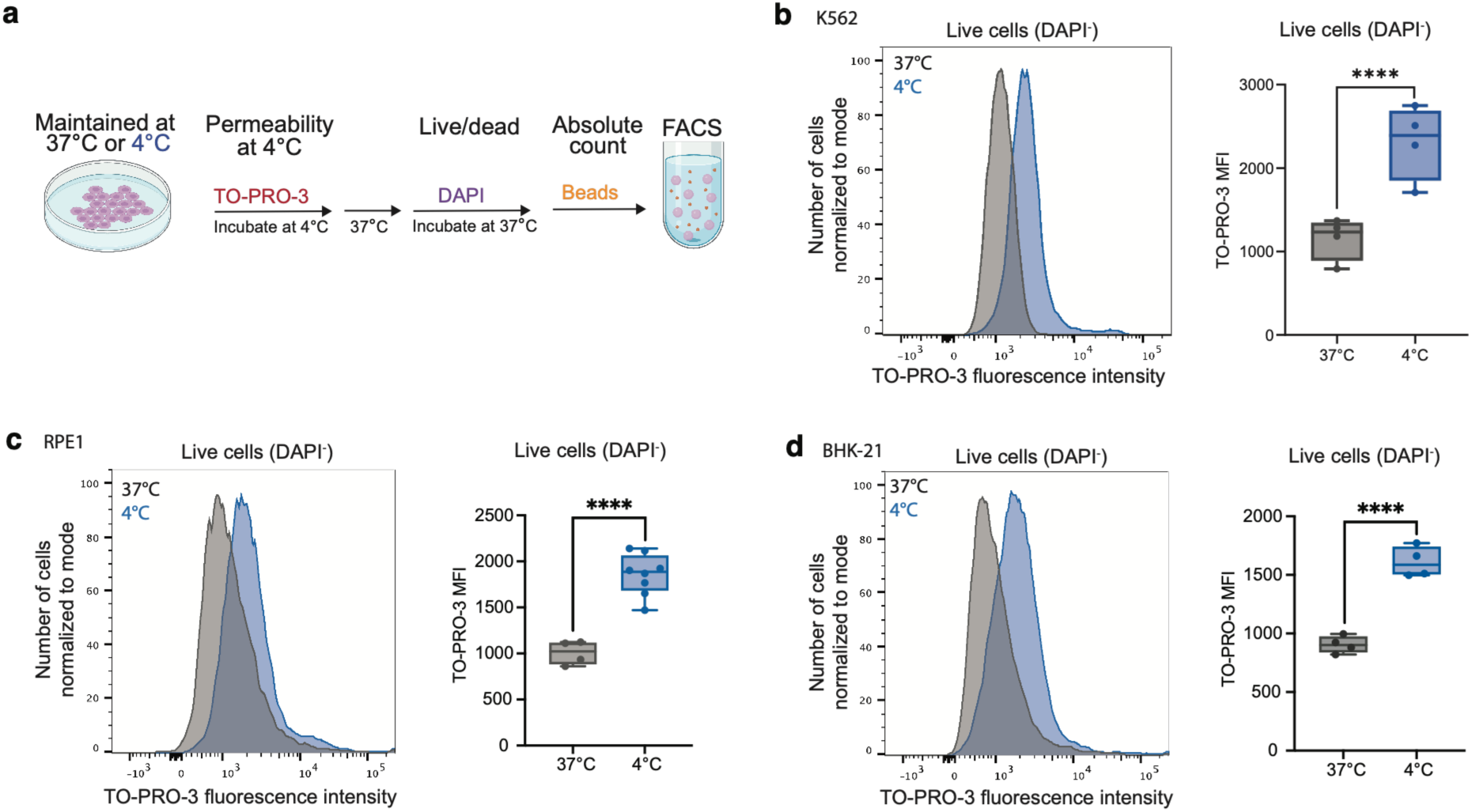
Assessment of plasma membrane permeability using a FACS-based assay. **a**, Schematic of multi-parameter flow cytometry assay to assess cell permeability in live cells. Cells were maintained for 1 day at 4°C or 37°C. To assess membrane permeability, cells were exposed for 15 minutes to the cell-impermeable dye TO-PRO-3 at 4°C and washed. To distinguish live from dead cells, cells were rewarmed to 37°C for 30 minutes and then exposed to the cell-impermeable dye DAPI. A known quantity of CountBright™ fluorescent beads was added to the sample and evenly mixed to enable determination of absolute cell counts during FACS**. b-d,** Representative plots of TO-PRO-3 fluorescent intensity for K562 (**b**), RPE1 (**c**), and BHK-21 (**d**) cells after 1 day at 4°C (blue) or 37°C (gray). Only live (DAPI negative) cells are plotted. TO-PRO-3 median fluorescence intensity (MFI) is plotted on the right (n = 4 samples for each condition). All values show mean ± SEM, with significance measured by student’s t-test. ****<0.0001.

**Supplement 3.**
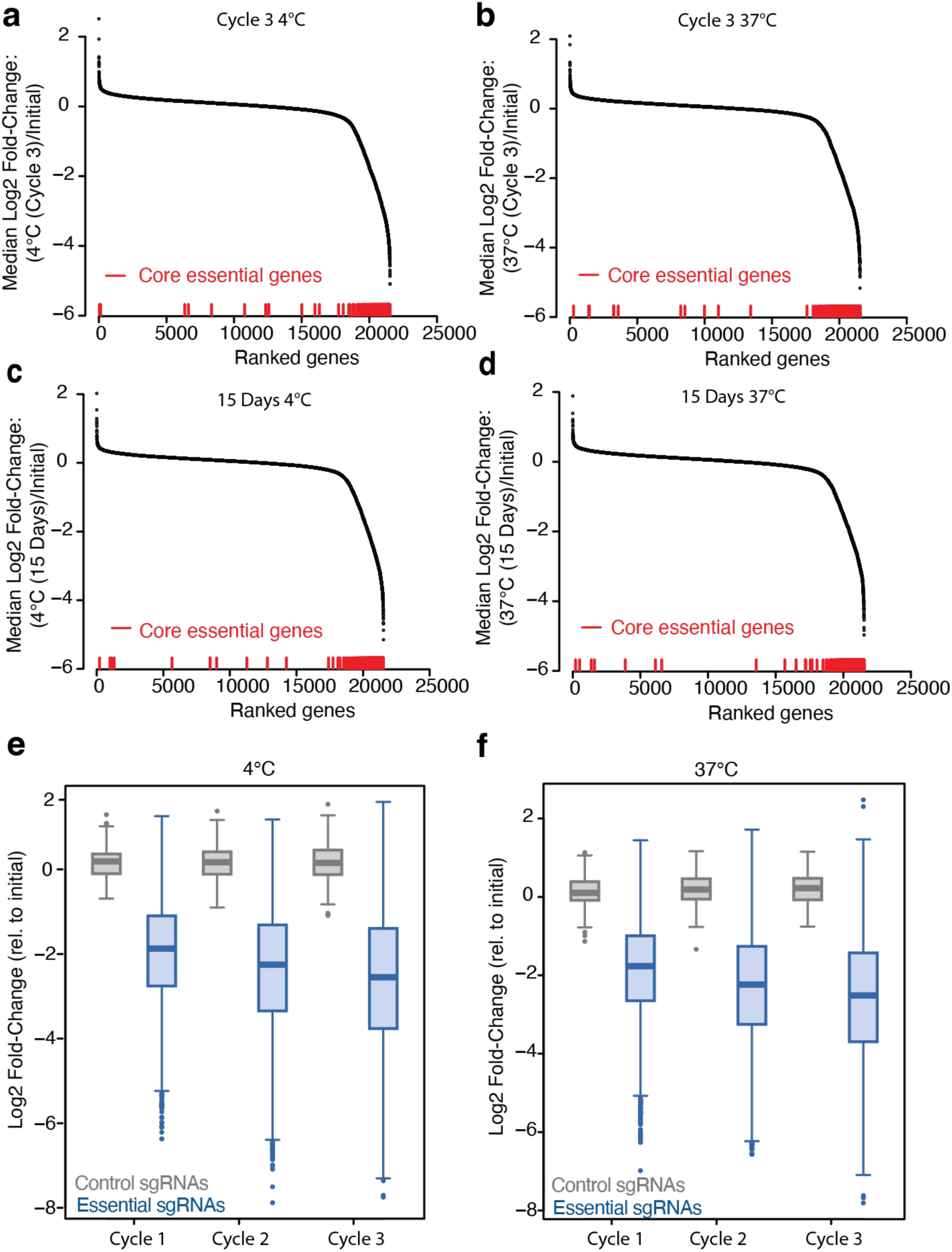
Depletion of Core Essential Genes in Genome-Wide BHK-21 Screens. **a-d**, Genes ranked by median fold-change (log2) in genome-wide BHK-21 screens. **a**, after three cycles of cold exposure and rewarming (Cycle 3 4°C), **b**, matched constant 37°C control condition (Cycle 3 37°C), **c**, after 15 days of 4°C exposure (15 Days 4°C), **d**, matched constant 37°C control condition (15 Days 37°C). Core essential genes^26^ indicated in red are positioned below based on gene rank to demonstrate their depletion in each screen condition. **e-f**, Boxplots showing log2 fold change in representation for the population of control sgRNAs (gray; n = 250) or sgRNAs targeting core essential genes^26^ (blue; n = 4635) over **e)** three cycles of cold exposure and rewarming (4°C) or **f)** constant 37°C control conditions. The line within each box represents the median, the bounds of each box represent the first and third quartiles, and the whiskers extend to the furthest data point within 1.5 times the interquartile range. A two-sided Kolmogorov-Smirnov test was used to test the difference between each pair of control/essential-gene-targeting sgRNA distributions (estimated p-value < 2.2e-16 for all six pairs in e and f).

**Supplement 4.**
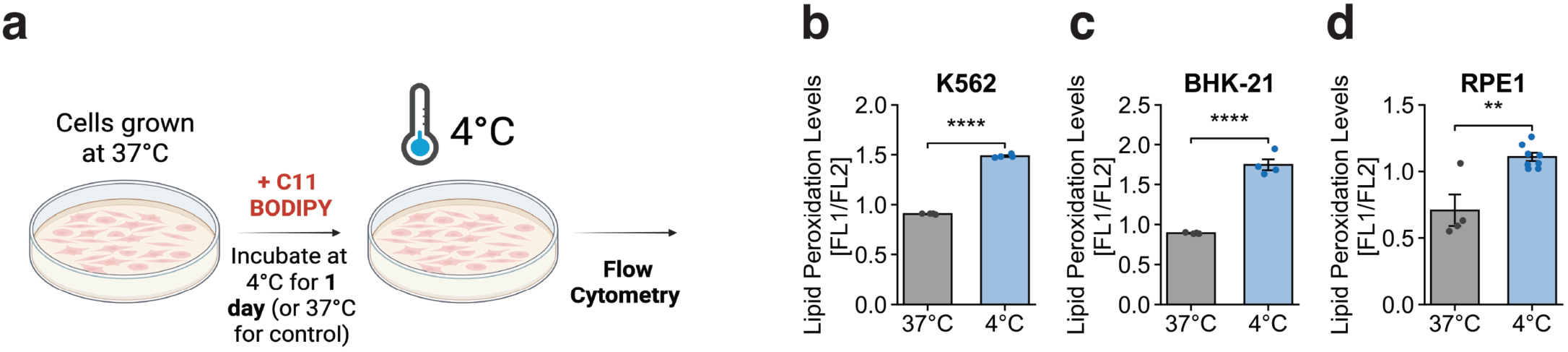
Cold exposure induces lipid peroxidation. **a**, Schematic of C11-BIODIPY (lipophilic ratiometric dye) loading into cells prior to exposure for 1 day to 37°C or 4°C, and flow cytometry. **b–d,** Lipid peroxidation, estimated as the ratio of the oxidized (green, FITC) to reduced (red, PE–Texas Red) forms of C11-BODIPY, in **(b)** K562, **(c)** BHK-21, and **(d)** RPE1 cells after 1 day at the indicated temperatures. Cold exposure increased the oxidized:reduced ratio in all three lines: K562 (37 °C vs 4 °C; n = 4 each), BHK-21 (37 °C vs 4 °C; n = 4 each), and RPE1 (37 °C, n = 4; 4 °C, n = 8). Bars show mean ± SEM with individual replicates overlaid; significance was assessed by unpaired two-tailed student’s t-test. **P < 0.01; ****P < 0.0001.

**Supplement 5.**
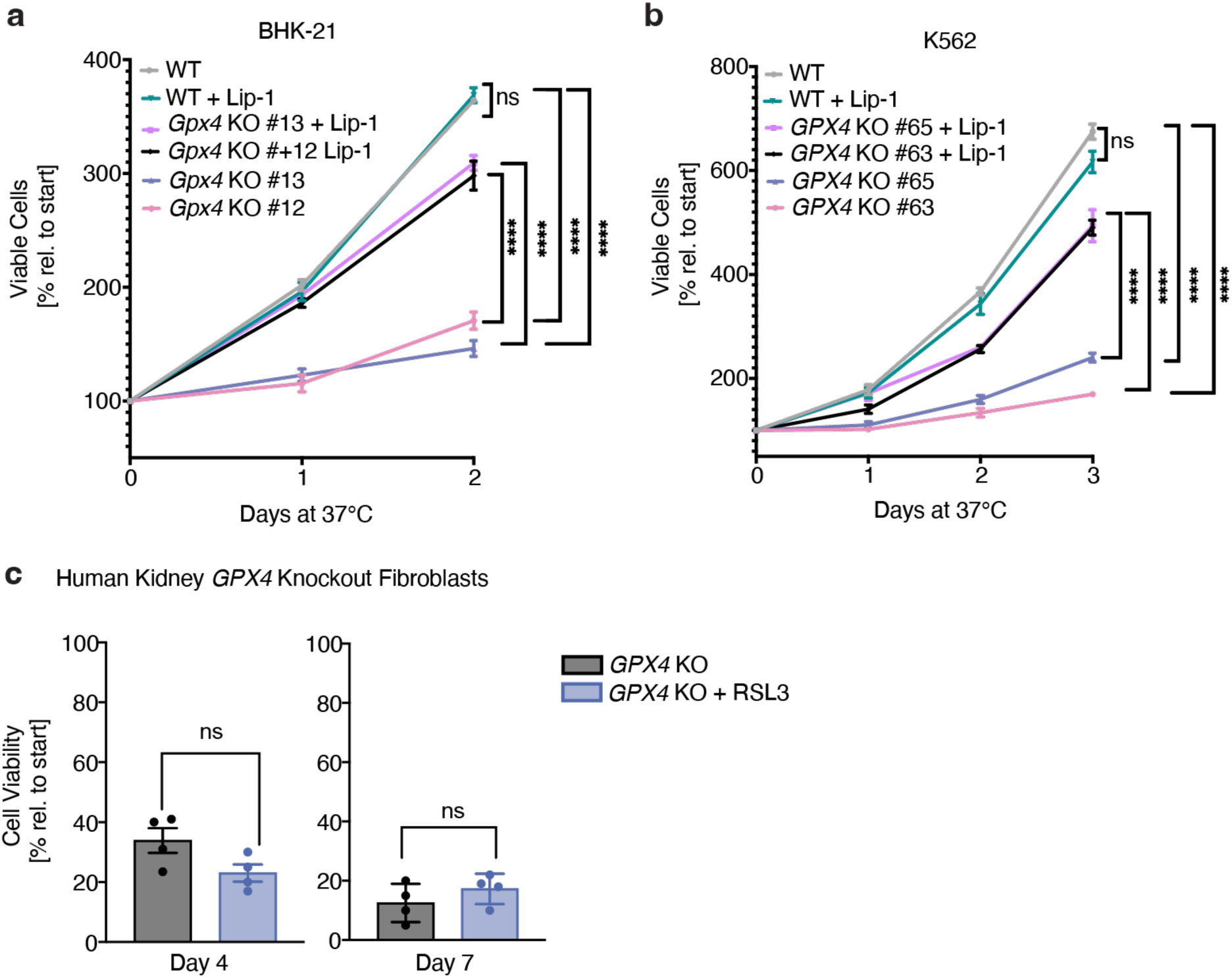
Growth of BHK21 and K562 *GPX4* knockout cells and effects of RSL3 treatment on human kidney fibroblast *GPX4* knockout cells. **a**, Viable K562 cells recorded as percentage relative to start based on trypan blue staining over the course of 3 days at 37°C. Cell growth is significantly decreased in *GPX4* K562 KO clones compared to WT K562 cells. Supplementation of liproxstatin-1 (2.5 µM) increases cell viability in *GPX4* KO cells and not WT cells at 37°C. **b**, Viable BHK-21 cells recorded as percentage relative to start based on trypan blue staining over the course of 2 days. Cell growth is significantly decreased in *Gpx4* BHK-21 KO cells compared to WT BHK-21 cells. Supplementation of liproxstatin-1 (2.5 µM) increases cell viability in *Gpx4* KO cell and not WT cells at 37°C. **c**, Human kidney GPX4 KO cells were placed at 4°C and left untreated or treated with RSL3 (1 μM) for 4, and 7 days (n = 4). Treatment with RSL3 does not confer significant additional death (n=4 per timepoint and condition) as measure by two-tailed t-test. All values show mean ± SEM, with significance measured by one-way ANOVA adjusted for multiple comparisons with Tukey’s HSD unless otherwise specified. \**P* < 0.05; \*\**P* < 0.01; \*\*\**P* < 0.001; \*\*\*\**P* < 0.0001; ns *P* > 0.05.

**Supplement 6.**
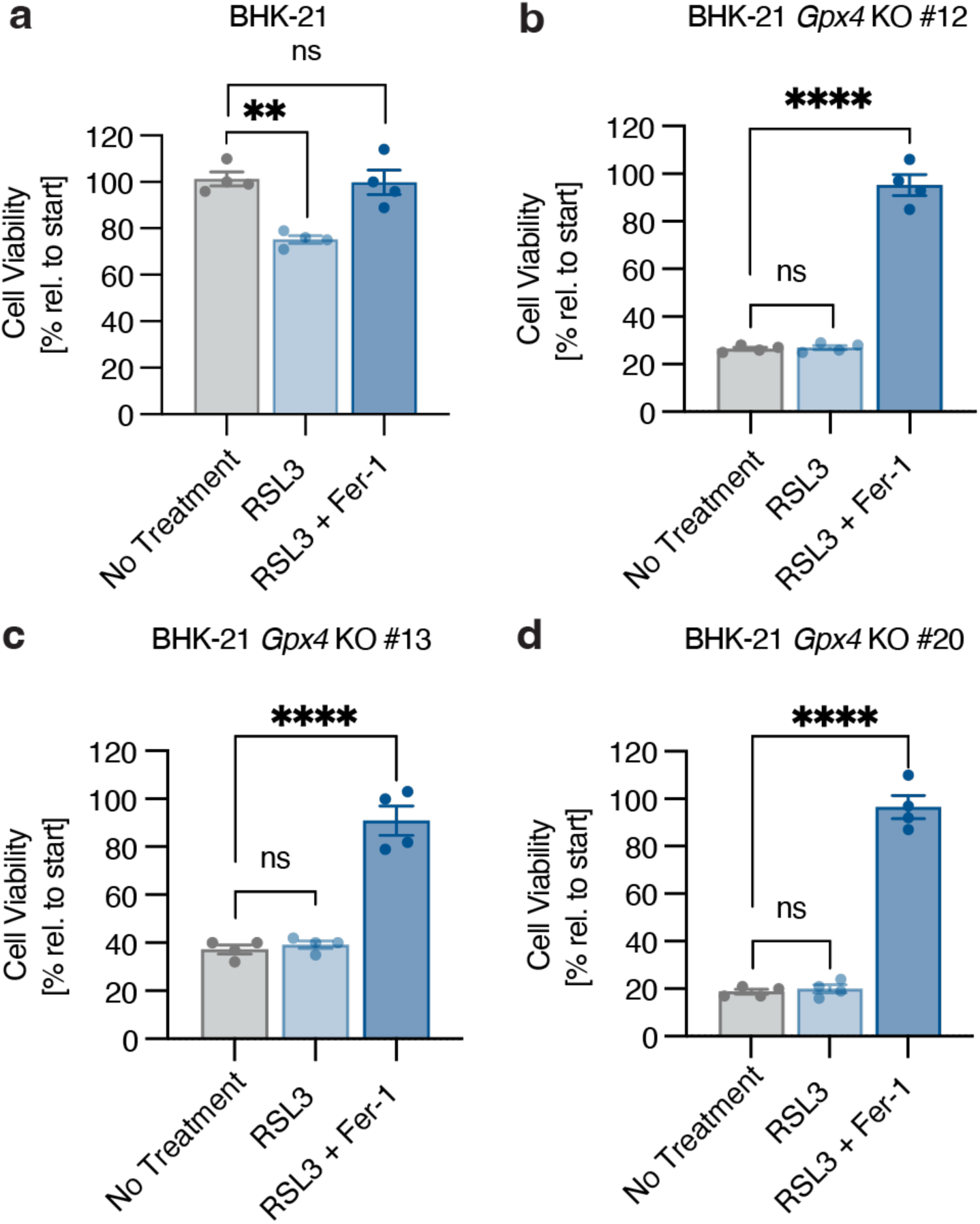
RSL3 treatment has no effect on the viability of cold-exposed *Gpx4* KO BHK-21 cells. **a-d**, Wild-type BHK-21 cells **(a)** and three independent *Gpx4* KO BHK-21 clonal lines **(b-d)** were treated with RSL3 (1 µM) and ferrostatin-1 (Fer-1, 1 µM) as indicated for 24 hours at 4°C (*n* = 4) prior to trypan blue staining. Wild-type BHK-21 cells show a significant decrease in cell viability when treated with RSL3 compared to no treatment (\*\**P* = 0.0013), whereas *Gpx4* KO lines show no significant changes in viability (ns, *P* > 0.05). All values show mean ± SEM, with significance measured by one-way ANOVA adjusted for multiple comparisons by Dunnett’s test. \**P* < 0.05; \*\**P* < 0.01; \*\*\**P* < 0.001; \*\*\*\**P* < 0.0001; ns *P* > 0.05.

**Supplement 7.**
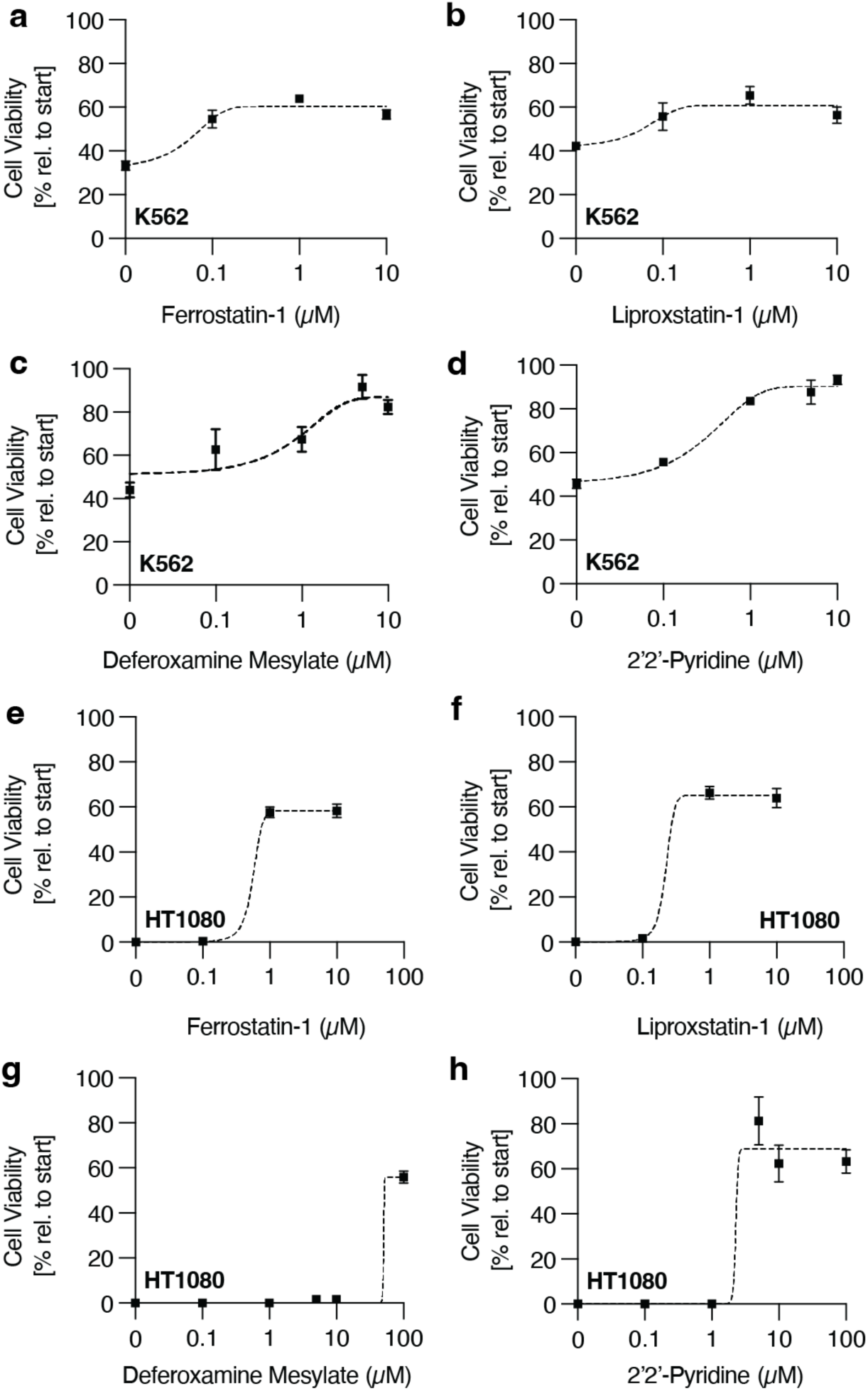
Ferroptosis inhibitors and iron chelators increase cold cell viability in a dose-dependent manner. **a-h**, K562 **(a-d)** and HT1080 **(e-h)** cells were treated with varying concentrations of the ferroptosis inhibitors, ferrostatin-1 and liproxstatin-1, and iron chelators, deferoxamine and 2’2’-pyridine, for four days at 4°C prior to assaying cell viability by trypan blue staining (*n* = 3). All values show mean ± SEM.

**Supplement 8.**
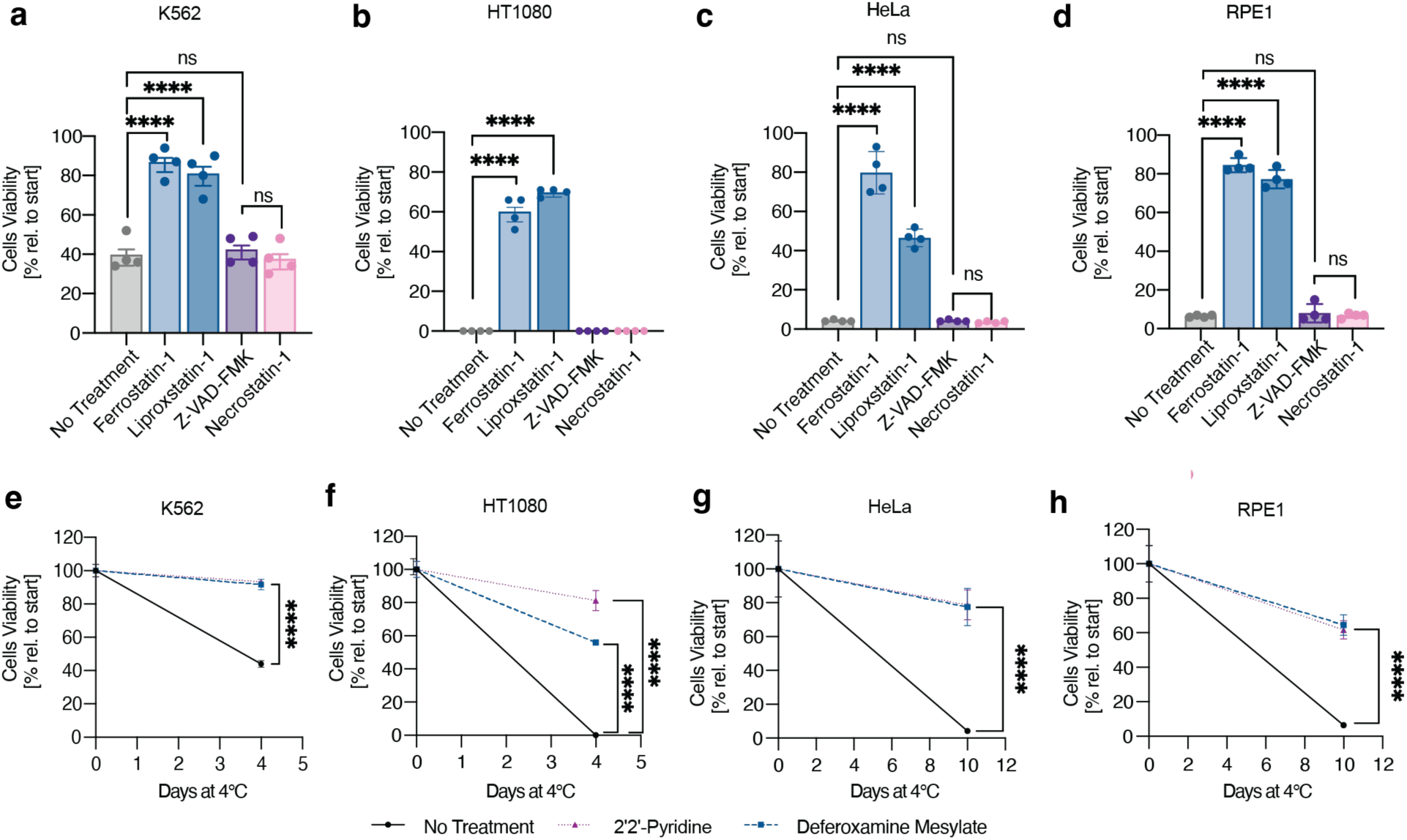
Cells derived from non-hibernators undergo cold-induced ferroptotic cell death. **a**, K562 cells were treated with ferrostatin-1 (1 µM) (\*\*\*\**P* < 0.0001), liproxstatin-1 (1 µM) (\*\*\*\**P* < 0.0001), Z-VAD-FMK (1 µM) (ns, *P* = 0.9760), or necrostatin-1 (1 µM) (ns, *P* = 0.9835) for 4 days at 4°C prior to trypan blue staining (*n* = 4). **b**, HT1080 cells treated with ferrostatin-1 (1 µM) (\*\*\*\**P* < 0.0001), liproxstatin-1 (1 µM) (\*\*\*\**P* < 0.0001), Z-VAD-FMK (1 µM), or necrostatin-1 (1 µM) for 4 days at 4°C prior to trypan blue staining (*n* = 4). **c**, HeLa cells treated with ferrostatin-1 (1 µM) (\*\*\*\**P* < 0.0001), liproxstatin-1 (1 µM) (\*\*\*\**P* < 0.0001), Z-VAD-FMK (1 µM) (ns, *P* > 0.9999), or necrostatin-1 (1 µM) (ns, *P* = 0.9987) for 10 days at 4°C prior to trypan blue staining (*n* = 4). **d**, RPE1 cells treated with ferrostatin-1 (1 µM) (\*\*\*\**P* < 0.0001), liproxstatin-1 (1 µM) (\*\*\*\**P* < 0.0001), Z-VAD-FMK (1 µM) (ns, *P* = 0.9291), or necrostatin-1 (1 µM) (ns, *P* > 0.9999) for 10 days at 4°C prior to trypan blue staining (*n* = 4). **e**, K562 cells treated with deferoxamine mesylate (5 µM) (\*\*\**P* = 0.0002) or 2’2’-pyridine (10 µM) (\*\*\*\**P* < 0.0001) for 4 days at 4°C prior to trypan blue staining (*n* = 3). **f**, HT1080 cells treated with deferoxamine mesylate (100 µM) (\*\*\*\**P* < 0.0001) or 2’2’-pyridine (5 µM) (\*\*\**P* = 0.0002) for 4 days at 4°C prior to trypan blue staining (*n* = 3). **g**, HeLa cells treated with deferoxamine mesylate (5 µM) (\*\*\*\**P* < 0.0001) or 2’2’-pyridine (5 µM) (\*\*\*\**P* < 0.0001) for 10 days at 4°C prior to trypan blue staining (*n* = 3). **h**, RPE1 cells treated with deferoxamine mesylate (100 µM) (\*\*\*\**P* < 0.0001) or 2’2’-pyridine (5 µM) (\*\*\**P* < 0.0001) for 10 days at 4°C prior to trypan blue staining (*n* = 3). All values show mean ± SEM, with significance measured one-way ANOVA adjusted for multiple comparisons by Dunnett’s test. \**P* < 0.05; \*\**P* < 0.01; \*\*\**P* < 0.001; \*\*\*\**P* < 0.0001; ns *P* > 0.05.

**Supplement 9.**
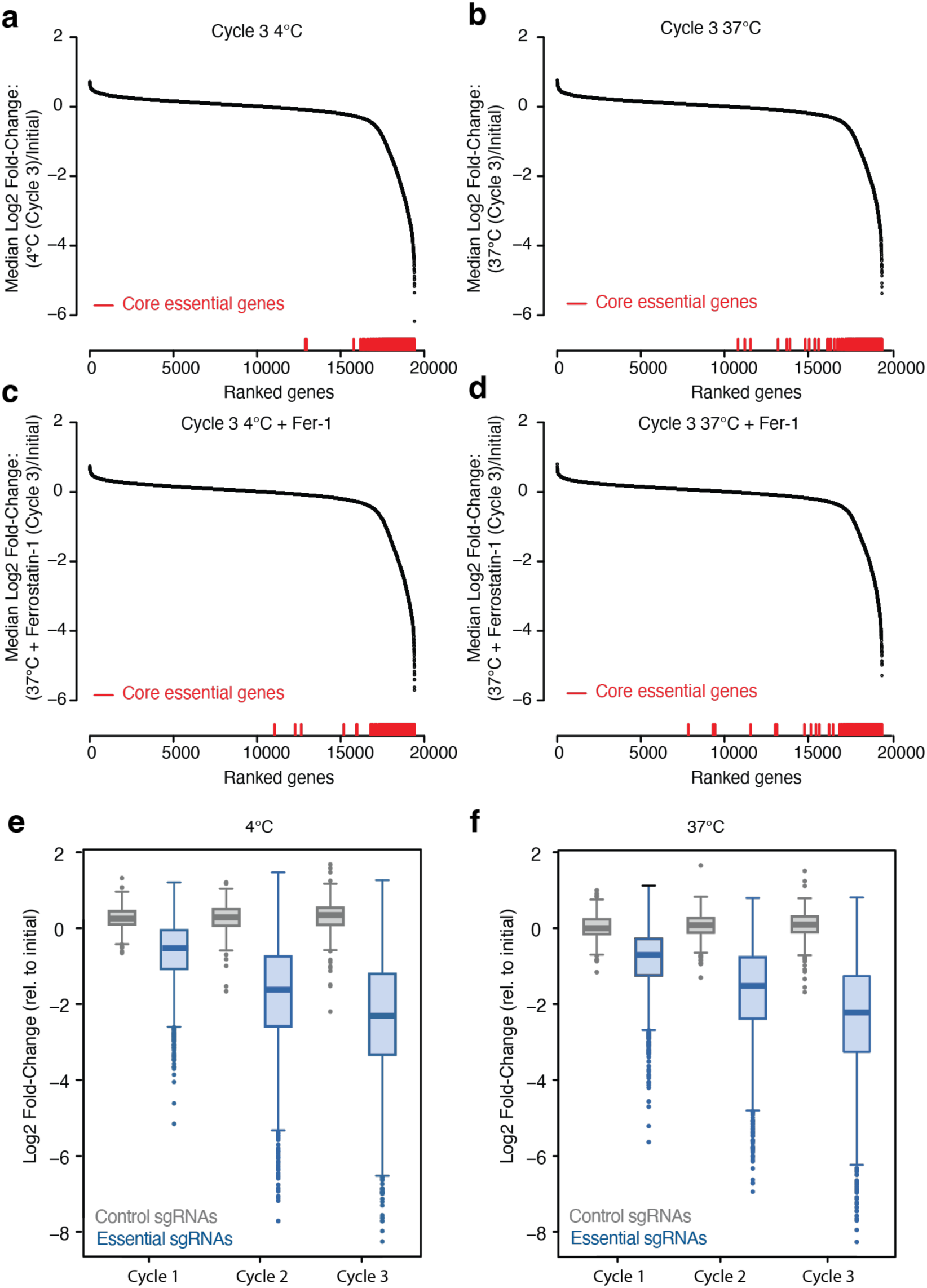
Depletion of Core Essential Genes in Genome-Wide K562 Screens. **a-d**, Genes ranked by median fold-change (log2) in genome-wide K562 screens. **a**, after three cycles of cold exposure (Cycle 3 4°C), **b**, matched constant 37°C control condition (Cycle 3 37°C), **c**, three cycles of cold exposure with 1 µM ferrostatin-1 (Cycle 3 4°C + Fer-1), **d**, matched constant 37°C control condition with 1 µM ferrostatin-1 (Cycle 3 37°C + Fer-1). Core essential genes^26^ (red bars) are positioned below based on gene rank to demonstrate their depletion in each screen condition. **e-f**, Boxplots showing the log2 fold change in representation for the population of control sgRNAs (gray; n = 500) or sgRNAs targeting core essential genes^26^ (blue; n = 3219) over (**e**) three cycles of cold exposure (4°C) or (**f**) constant 37°C control conditions. The line within each box represents the median, the bounds of each box represent the first and third quartiles, and the whiskers extend to the furthest data point within 1.5 times the interquartile range. A two-sided Kolmogorov-Smirnov test was used to test the difference between each pair of control/essential-gene-targeting sgRNA distributions (estimated p-value < 2.2e-16 for all six pairs in e and f).

**Supplement 10.**
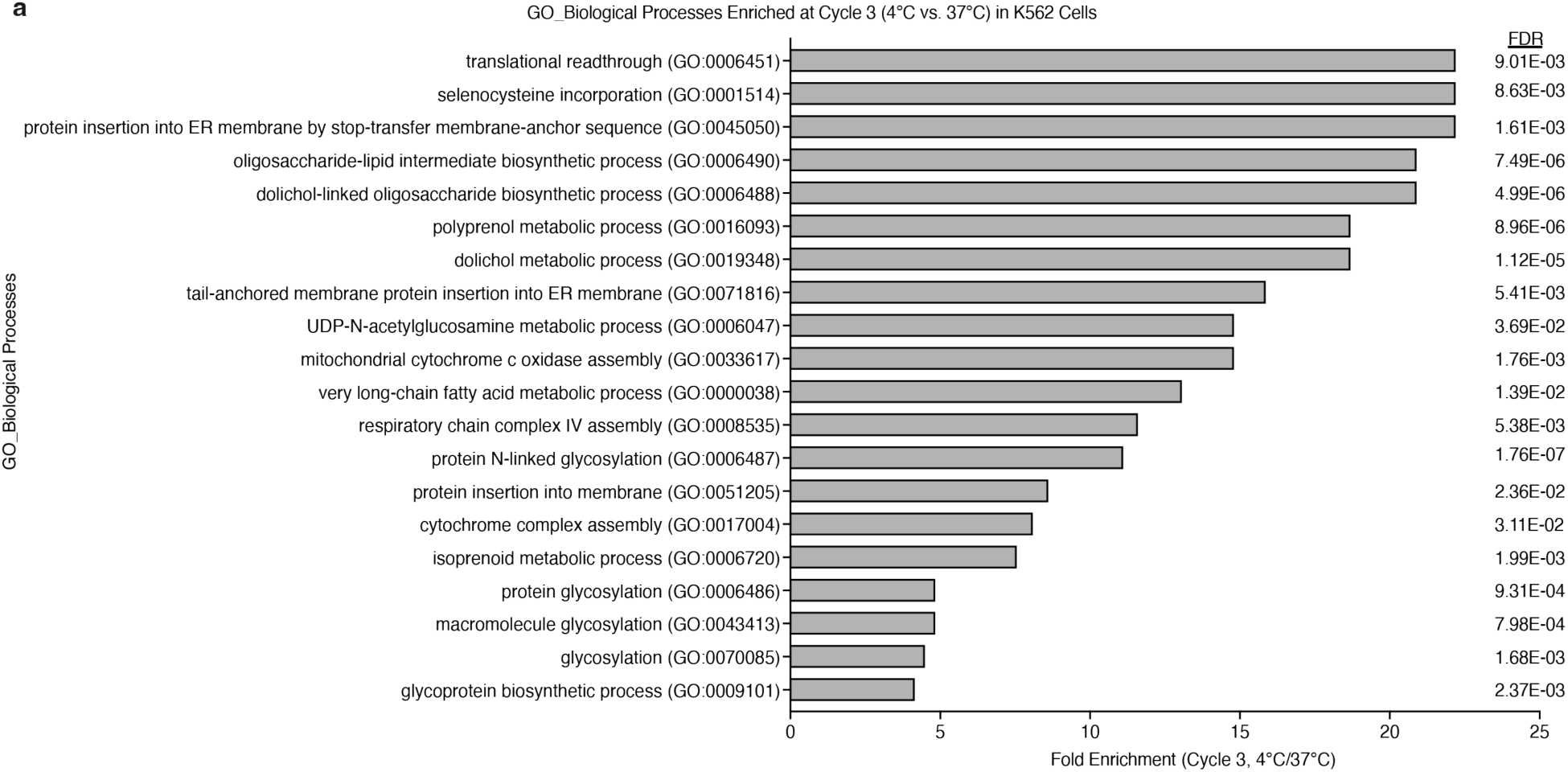
Top 20 enriched pathways at Cycle 3 (4°C vs. 37°C) in K562 cells. **a**, Graphical representation of the top 20 enriched, differentially expressed gene sets (204 genes; FDR < 0.1) from the genome-scale CRISPR-Cas9 screen in K562 cells (Cycle 3 4°C vs. 3 passages at 37°C). GO_Biological Processes Panther overrepresentation test was used to determine enriched gene sets. Functional annotation analysis of the selectively required genes identified pathways related to translational readthrough, selenocysteine incorporation, protein insertion into the ER, glycosylation, fatty acid metabolism, and mitochondrial respiration. A full list of pathways is provided in Table S6.

**Supplement 11.**
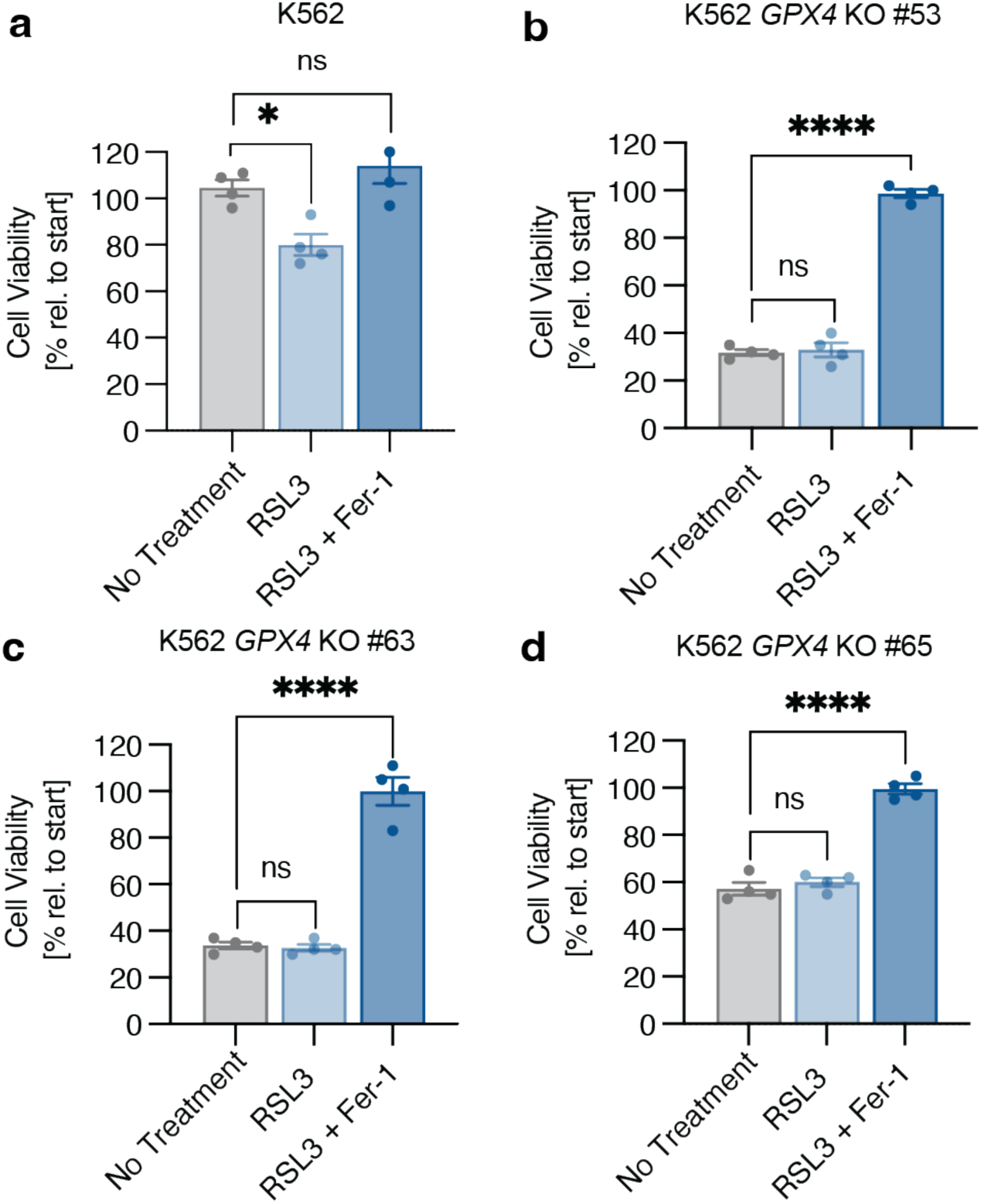
RSL3 treatment has no effect on the viability of cold-exposed *GPX4* KO K562 cells. **a-d**, Wild-type K562 cells **(a)** and three independent *GPX4* KO K562 clonal lines **(b-d)** were treated with RSL3 (1 µM) and ferrostatin-1 (Fer-1, 1 µM) as indicated for 8 hours at 4°C (*n* = 4) prior to trypan blue staining. Wild-type K562 cells show a significant decrease in cell viability when treated with RSL3 compared to no treatment (\**P* = 0.0213), whereas *GPX4* KO lines show no significant changes in viability (ns, *P* > 0.05). All values show mean ± SEM, with significance measured by one-way ANOVA adjusted for multiple comparisons by Dunnett’s test. \**P* < 0.05; \*\**P* < 0.01; \*\*\**P* < 0.001; \*\*\*\**P* < 0.0001; ns *P* > 0.05.

**Supplement 12.**
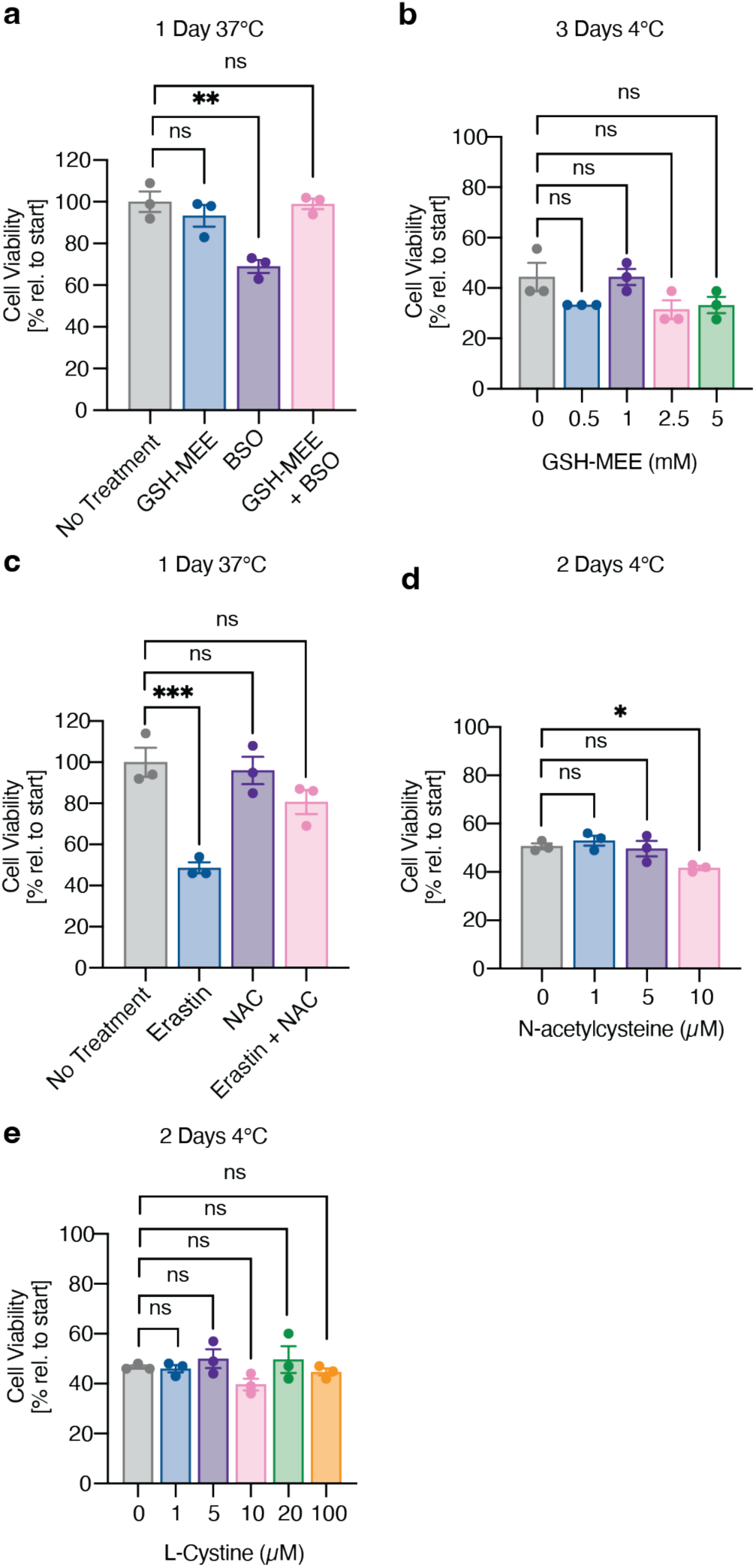
Cell permeable glutathione and glutathione precursors do not increase cold cell viability. **a,** K562 cells were placed at 37°C and treated with cell permeable glutathione GSH-MEE (1 µM) and/or the glutathione synthesis inhibitor buthionine sulfoximine (BSO, 1 µM) for 1 day (*n* = 3). Treatment with BSO resulted in increased cell death (\*\**P* = 0.0018) that was rescued by GSH-MEE (ns, *P* = 0.9961). **b,** Treatment with GSH-MEE has no effect on K562 cold-induced death after 3 days at 4°C as measured by trypan blue staining (*n* = 3, 5 µM; ns, *P* = 0.1558). **c,** K562 cells were placed at 37°C and treated with ferroptosis inducer Erastin (10 µM) and N-acetylcysteine (NAC, 10 µM) for 1 day (*n* = 3). Treatment with Erastin resulted in increased cell death (\*\*\**P* = 0.0006) that was rescued by NAC (ns, *P* = 0.1091). **d,** Treatment with N-acetylcysteine does not increase cold cell viability in K562 cells after 2 days at 4°C as measured by trypan blue staining (*n* = 3, 10 µM; \**P* = 0.0352). **e,** Treatment with L-Cystine does not increase cold cell viability in K562 cells after 2 days at 4°C as measured by trypan blue staining (*n* = 3, 100 µM; ns, *P* = 0.9843). All values show mean ± SEM, with significance measured by one-way ANOVA adjusted for multiple comparisons by Dunnett’s test. \**P* < 0.05; \*\**P* < 0.01; \*\*\**P* < 0.001; \*\*\*\**P* < 0.0001; ns *P* > 0.05.

**Supplement 13.**
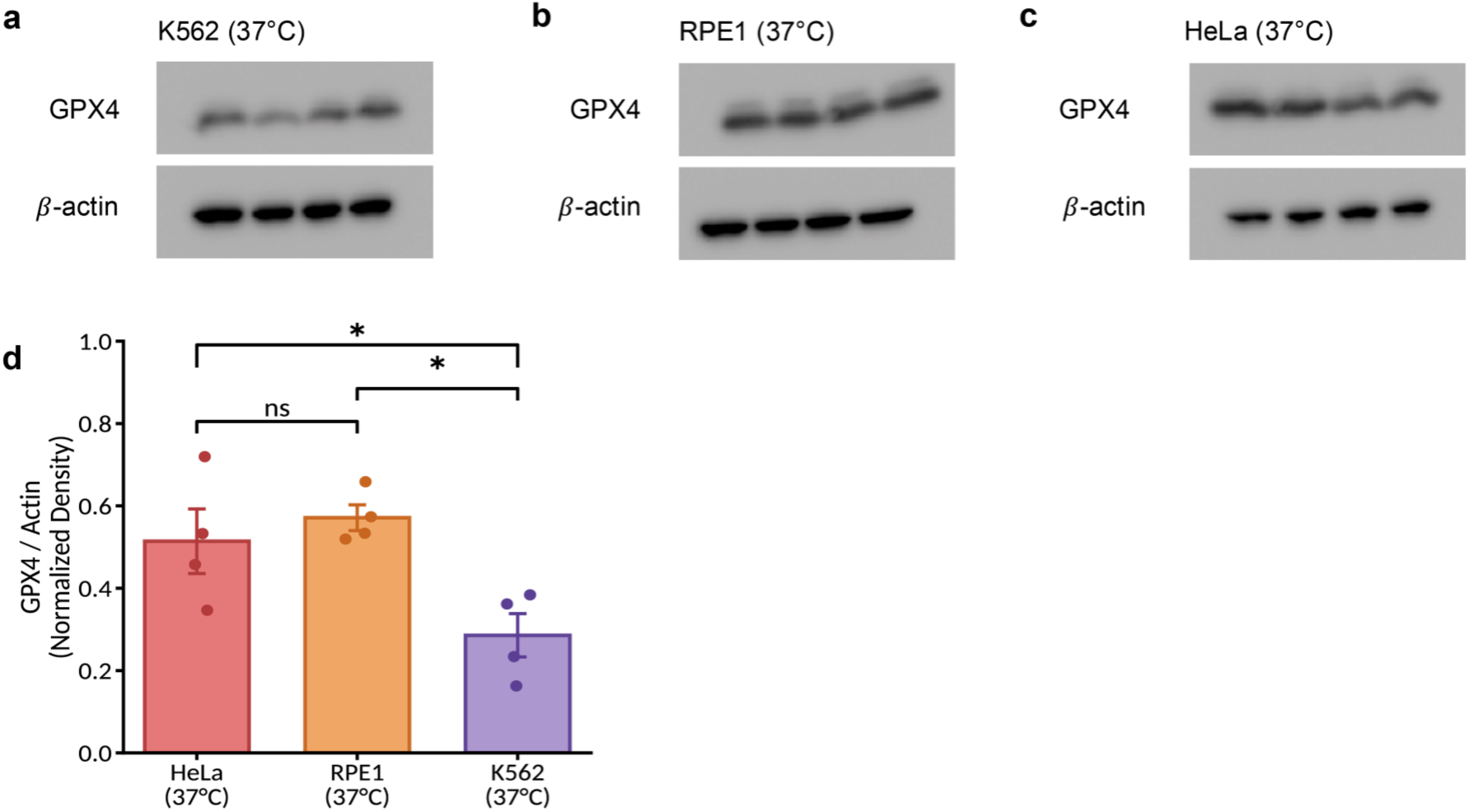
GPX4 protein levels across human cell lines. **a-c,** Western blotting for GPX4 and β-actin (loading control, n = 4 samples per cell line) in K562 (**a**), RPE1 (**b**) and HeLa (**c**) cell lines cultured at 37°C (24 hours following seeding). **d,** Quantified GPX4 immunoblot signal normalized to β-actin from **a-c** (n = 4 independent replicates for each). All values show mean ± SEM, with significance measured by one-way ANOVA followed by Tukey’s HSD post hoc test for all pairwise comparisons. *p < 0.05; ns, not significant. Densitometry was conducted using ImageJ.

**Supplement 14.**
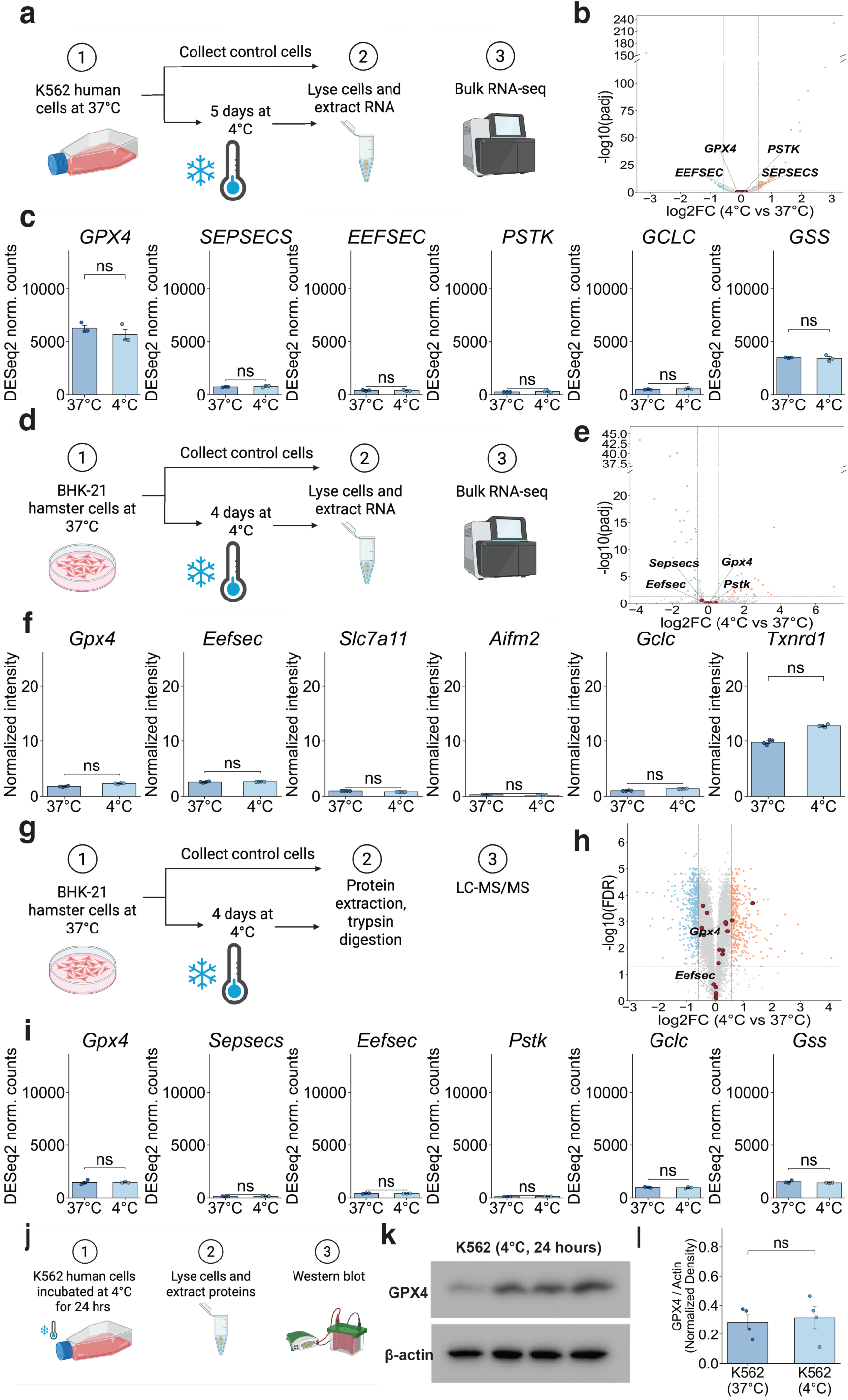
GPX4 mRNA and protein levels following exposure to 4°C. **a-c,** Bulk RNA-seq of K562 cells after 5 days of culture at 4°C versus standard 37°C conditions (n = 3). **a,** Schematic of the experimental design. **b,** Volcano plot. **c,** Selected GPX4 and ferroptosis/selenocysteine-pathway genes from (**b**). **d-f,** Bulk RNA-seq of BHK-21 cells after 4 days of culture at 4°C versus standard 37°C conditions (n = 3). **d,** Schematic of the experimental design. **e,** Volcano plot. **f**, Selected GPX4 and ferroptosis/selenocysteine-pathway gene from (**e**). **g-i,** Proteomics (TMT) of BHK-21 cells after 4 days of culture at 4°C versus standard 37°C conditions (n = 4). **g,** Schematic of the experimental design. **h,** Volcano plot. **i,** Selected GPX4 and ferroptosis/selenocysteine-pathway proteins from (**h**). Levels of Sepsecs and Pstk were below the level of detection by MS and are not shown. All values show mean ± SEM. **j-l,** Western blotting for GPX4 and β-actin (n = 4 samples per condition) in K562 cells cultured at 4°C for 24 hours. **j,** Schematic of the experimental design. **k,** Western blot. **l,** Quantified GPX4 immunoblot signal normalized to β-actin (n = 4 independent replicates per condition). Densitometry was conducted using ImageJ. All values show mean ± SEM. For **a-i,** genes/proteins were considered significantly changed if they passed both the test noted above (Benjamini-Hochberg FDR < 0.05) and an absolute log2 fold-change > 0.58 (1.5-fold); *FDR < 0.05; **FDR < 0.01; ***FDR < 0.001; ns, did not meet both criteria. For **k**, significance was measured by Welch’s two-tailed t-test; *p < 0.05; **p < 0.01; ***p < 0.001; ns, not significant.

**Supplement 15.**
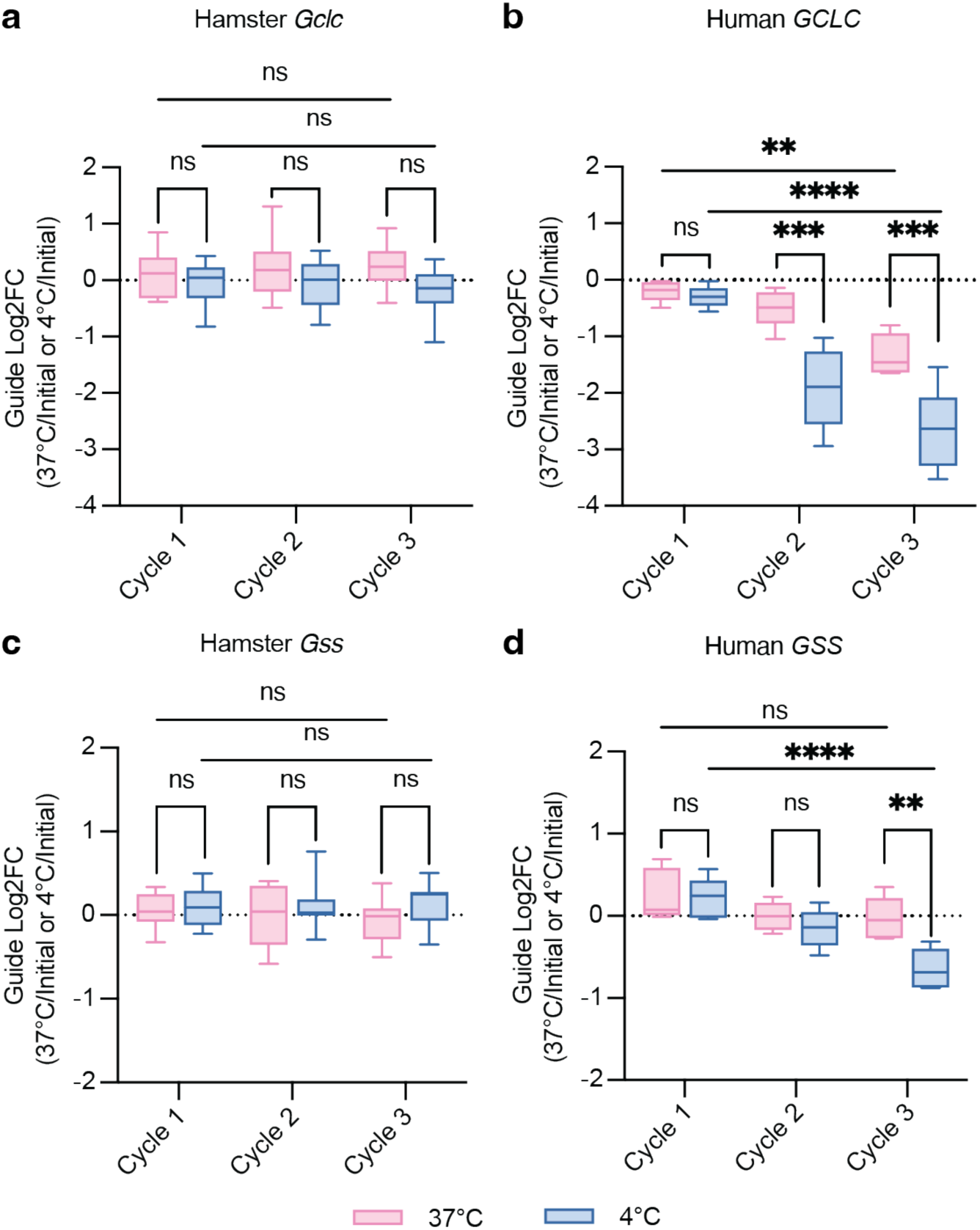
Glutathione biosynthesis genes show increased depletion in cold-exposed K562 cells. **a,c**, Log2 fold-change (Log2FC) of 10 guides per targeted gene in BHK-21 genome-wide screen, showing guide depletion over three cycles of cold exposure and rewarming. **a**, *Gclc*, **c**, *Gss*. **b**, **d**, Log2 fold-change (Log2FC) of 5 guides per targeted gene in K562 genome-wide screen, showing significant guide depletion over three cycles of cold exposure. **b**, *GCLC*, **d**, *GSS*. Significance between Cycle 1 versus Cycle 3 is measured by two-way ANOVA adjusted for multiple comparisons by Dunnett’s test. Significance between 37°C and 4°C for each cycle is measured by two-way ANOVA adjusted for multiple comparisons by Bonferroni’s test. \**P* < 0.05; \*\**P* < 0.01; \*\*\**P* < 0.001; \*\*\*\**P* < 0.0001; ns *P* > 0.05.

## References

1. Cheshire, W.P. (2016). Thermoregulatory disorders and illness related to heat and cold stress. Autonomic Neuroscience 196, 91–104. 10.1016/j.autneu.2016.01.001.

2. Physiology, Thermal Regulation - StatPearls - NCBI Bookshelf https://www.ncbi.nlm.nih.gov/books/NBK499843/.

3. Tan, C.L., and Knight, Z.A. (2018). Regulation of body temperature by the nervous system. Neuron 98, 31–48. 10.1016/j.neuron.2018.02.022.

4. Morrison, S.F., and Nakamura, K. (2019). Central Mechanisms for Thermoregulation. Annu Rev Physiol 81, 285–308. 10.1146/annurev-physiol-020518-114546.

5. Hendriks, K.D.W., Joschko, C.P., Hoogstra-Berends, F., Heegsma, J., Faber, K.-N., and Henning, R.H. (2020). Hibernator-Derived Cells Show Superior Protection and Survival in Hypothermia Compared to Non-Hibernator Cells. IJMS 21, 1864. 10.3390/ijms21051864.

6. Ou, J., Ball, J.M., Luan, Y., Zhao, T., Miyagishima, K.J., Xu, Y., Zhou, H., Chen, J., Merriman, D.K., Xie, Z., et al. (2018). iPSCs from a Hibernator Provide a Platform for Studying Cold Adaptation and Its Potential Medical Applications. Cell 173, 851–863.e16. 10.1016/j.cell.2018.03.010.

7. Hendriks, K.D.W., Lupi, E., Hardenberg, M.C., Hoogstra-Berends, F., Deelman, L.E., and Henning, R.H. (2017). Differences in mitochondrial function and morphology during cooling and rewarming between hibernator and non-hibernator derived kidney epithelial cells. Sci Rep 7, 15482. 10.1038/s41598-017-15606-z

8. Anegawa, D., Sugiura, Y., Matsuoka, Y., Sone, M., Shichiri, M., Otsuka, R., Ishida, N., Yamada, K., Suematsu, M., Miura, M., et al. (2021). Hepatic resistance to cold ferroptosis in a mammalian hibernator Syrian hamster depends on effective storage of diet-derived α-tocopherol. Commun Biol 4, 796. 10.1038/s42003-021-02297-6.

9. Park, H.G., Han, S.I., Oh, S.Y., and Kang, H.S. (2005). Cellular responses to mild heat stress. CMLS, Cell. Mol. Life Sci. 62, 10–23. 10.1007/s00018-004-4208-7.

10. Richter, K., Haslbeck, M., and Buchner, J. (2010). The Heat Shock Response: Life on the Verge of Death. Molecular Cell 40, 253–266. 10.1016/j.molcel.2010.10.006.

11. Karunanithi, S., and Brown, I.R. (2015). Heat shock response and homeostatic plasticity. Front. Cell. Neurosci. 9. 10.3389/fncel.2015.00068.

12. Kruuv, J., Glofcheski, D., Cheng, K.H., Campbell, S.D., Al-Qysi, H.M., Nolan, W.T., and Lepock, J.R. (1983). Factors influencing survival and growth of mammalian cells exposed to hypothermia. I. Effects of temperature and membrane lipid perturbers. J Cell Physiol 115, 179–185. 10.1002/jcp.1041150212.

13. Hochachka, P.W. (1986). Defense strategies against hypoxia and hypothermia. Science 231, 234–241. 10.1126/science.2417316.

14. Rule, G.S., Frim, J., Thompson, J.E., Lepock, J.R., and Kruuv, J. (1978). The effect of membrane lipid perturbers on survival of mammalian cells to cold. Cryobiology 15, 408–414. 10.1016/0011-2240(78)90059-7.

15. Zachariassen, K.E. (1991). Hypothermia and cellular physiology. Arctic Med Res 50 *Suppl 6*, 13–17.

16. Heller, H.C., and Hammel, H.T. (1972). CNS control of body temperature during hibernation. Comp Biochem Physiol A Comp Physiol 41, 349–359. 10.1016/0300-9629(72)90066-7.

17. Sunagawa, G.A., and Takahashi, M. (2016). Hypometabolism during Daily Torpor in Mice is Dominated by Reduction in the Sensitivity of the Thermoregulatory System. Sci Rep 6, 37011. 10.1038/srep37011.

18. Geiser, F. (2004). Metabolic rate and body temperature reduction during hibernation and daily torpor. Annu Rev Physiol 66, 239–274. 10.1146/annurev.physiol.66.032102.115105.

19. Chayama, Y., Ando, L., Tamura, Y., Miura, M., and Yamaguchi, Y. (2016). Decreases in body temperature and body mass constitute pre-hibernation remodelling in the Syrian golden hamster, a facultative mammalian hibernator. R Soc Open Sci 3, 160002. 10.1098/rsos.160002.

20. Park, K.J., Jones, G., and Ransome, R.D. (2000). Torpor, arousal and activity of hibernating Greater Horseshoe Bats (Rhinolophus ferrumequinum). Functional Ecology 14, 580–588. 10.1046/j.1365-2435.2000.t01-1-00460.x.

21. Cooper, S.T., Richters, K.E., Melin, T.E., Liu, Z., Hordyk, P.J., Benrud, R.R., Geiser, L.R., Cash, S.E., Simon Shelley, C., Howard, D.R., et al. (2012). The hibernating 13-lined ground squirrel as a model organism for potential cold storage of platelets. Am J Physiol Regul Integr Comp Physiol 302, R1202–R1208. 10.1152/ajpregu.00018.2012.

22. Heldmaier, G., Ortmann, S., and Elvert, R. (2004). Natural hypometabolism during hibernation and daily torpor in mammals. Respiratory Physiology & Neurobiology 141, 317–329. 10.1016/j.resp.2004.03.014.

23. Zhang, X., Ge, L., Jin, G., Liu, Y., Yu, Q., Chen, W., Chen, L., Dong, T., Miyagishima, K.J., Shen, J., et al. (2024). Cold-induced FOXO1 nuclear transport aids cold survival and tissue storage. Nat Commun 15, 2859. 10.1038/s41467-024-47095-w.

24. Zhang, X., Chen, L., Liu, W., Shen, J., Sun, H., Liang, J., Lv, G., Chen, G., Yang, Y., and Ou, J. (2022). 5-Aminolevulinate improves metabolic recovery and cell survival of the liver following cold preservation. Theranostics 12, 2908–2927. 10.7150/thno.69446.

25. Harris, R.A., Raveendran, M., Lyfoung, D.T., Sedlazeck, F.J., Mahmoud, M., Prall, T.M., Karl, J.A., Doddapaneni, H., Meng, Q., Han, Y., et al. (2022). Construction of a new chromosome-scale, long-read reference genome assembly for the Syrian hamster, Mesocricetus auratus. Gigascience 11, giac039. 10.1093/gigascience/giac039.

26. Hart, T., Tong, A.H.Y., Chan, K., Van Leeuwen, J., Seetharaman, A., Aregger, M., Chandrashekhar, M., Hustedt, N., Seth, S., Noonan, A., et al. (2017). Evaluation and Design of Genome-Wide CRISPR/SpCas9 Knockout Screens. G3 (Bethesda) 7, 2719–2727. 10.1534/g3.117.041277.

27. Summer, S., Smirnova, A., Gabriele, A., Toth, U., Fasemore, A.M., Förstner, K.U., Kuhn, L., Chicher, J., Hammann, P., Mitulović, G., et al. (2020). YBEY is an essential biogenesis factor for mitochondrial ribosomes. Nucleic Acids Res 48, 9762–9786. 10.1093/nar/gkaa148.

28. Ni, M., Solmonson, A., Pan, C., Yang, C., Li, D., Notzon, A., Cai, L., Guevara, G., Zacharias, L.G., Faubert, B., et al. (2019). Functional Assessment of Lipoyltransferase-1 Deficiency in Cells, Mice, and Humans. Cell Rep 27, 1376–1386.e6. 10.1016/j.celrep.2019.04.005.

29. Xu, G., Li, W., Zhao, Y., Fan, T., Gao, Q., Wang, Y., Zhang, F., Gao, M., An, Z., and Yang, Z. (2024). Overexpression of Lias Gene Alleviates Cadmium-Induced Kidney Injury in Mice Involving Multiple Effects: Metabolism, Oxidative Stress, and Inflammation. Biol Trace Elem Res 202, 2797–2811. 10.1007/s12011-023-03883-x.

30. Douce, R., Bourguignon, J., Neuburger, M., and Rébeillé, F. (2001). The glycine decarboxylase system: a fascinating complex. Trends Plant Sci 6, 167–176. 10.1016/s1360-1385(01)01892-1.

31. Cronan, J.E. (2020). Progress in the Enzymology of the Mitochondrial Diseases of Lipoic Acid Requiring Enzymes. Front. Genet. 11, 510. 10.3389/fgene.2020.00510.

32. Weaver, K., and Skouta, R. (2022). The Selenoprotein Glutathione Peroxidase 4: From Molecular Mechanisms to Novel Therapeutic Opportunities. Biomedicines 10, 891. 10.3390/biomedicines10040891.

33. Ingold, I., Berndt, C., Schmitt, S., Doll, S., Poschmann, G., Buday, K., Roveri, A., Peng, X., Porto Freitas, F., Seibt, T., et al. (2018). Selenium Utilization by GPX4 Is Required to Prevent Hydroperoxide-Induced Ferroptosis. Cell 172, 409–422.e21. 10.1016/j.cell.2017.11.048.

34. Seibt, T.M., Proneth, B., and Conrad, M. (2019). Role of GPX4 in ferroptosis and its pharmacological implication. Free Radical Biology and Medicine 133, 144–152. 10.1016/j.freeradbiomed.2018.09.014.

35. Park, S., Park, M.J., Kwon, E.-J., Oh, J.-Y., Chu, Y.-J., Kim, H.S., Park, S., Kim, T.H., Kwon, S.W., Kim, Y.S., et al. (2025). The protective role of GPX4 in naïve ESCs is highlighted by induced ferroptosis resistance through GPX4 expression. Redox Biology 81, 103539. 10.1016/j.redox.2025.103539.

36. Seiler, A., Schneider, M., Förster, H., Roth, S., Wirth, E.K., Culmsee, C., Plesnila, N., Kremmer, E., Rådmark, O., Wurst, W., et al. (2008). Glutathione Peroxidase 4 Senses and Translates Oxidative Stress into 12/15-Lipoxygenase Dependent- and AIF-Mediated Cell Death. Cell Metabolism 8, 237–248. 10.1016/j.cmet.2008.07.005.

37. Varlamova, E.G., Goltyaev, M.V., Novoselov, S.V., Novoselov, V.I., and Fesenko, E.E. (2013). Selenocysteine biosynthesis and mechanism of incorporation into growing proteins. Mol Biol 47, 488–495. 10.1134/S0026893313040134.

38. French, R., and Simonovic, M. (2012). Synthesis and decoding of selenocysteine and human health. Croatian medical journal 53, 535–550. 10.3325/cmj.2012.53.535.

39. Carlson, B.A., Xu, X.-M., Kryukov, G.V., Rao, M., Berry, M.J., Gladyshev, V.N., and Hatfield, D.L. (2004). Identification and characterization of phosphoseryl-tRNA^[Ser]Sec^ kinase. Proc. Natl. Acad. Sci. U.S.A. 101, 12848–12853. 10.1073/pnas.0402636101.

40. Ganichkin, O.M., Xu, X.-M., Carlson, B.A., Mix, H., Hatfield, D.L., Gladyshev, V.N., and Wahl, M.C. (2008). Structure and Catalytic Mechanism of Eukaryotic Selenocysteine Synthase. Journal of Biological Chemistry 283, 5849–5865. 10.1074/jbc.M709342200.

41. Donovan, J., and Copeland, P.R. (2010). The Efficiency of Selenocysteine Incorporation Is Regulated by Translation Initiation Factors. Journal of Molecular Biology 400, 659–664. 10.1016/j.jmb.2010.05.026.

42. Gupta, N., DeMong, L.W., Banda, S., and Copeland, P.R. (2013). Reconstitution of Selenocysteine Incorporation Reveals Intrinsic Regulation by SECIS Elements. Journal of Molecular Biology 425, 2415–2422. 10.1016/j.jmb.2013.04.016.

43. Donovan, J., and Copeland, P.R. (2010). Threading the Needle: Getting Selenocysteine Into Proteins. Antioxidants & Redox Signaling 12, 881–892. 10.1089/ars.2009.2878.

44. Gonzalez-Flores, J.N., Gupta, N., DeMong, L.W., and Copeland, P.R. (2012). The Selenocysteine-specific Elongation Factor Contains a Novel and Multi-functional Domain. Journal of Biological Chemistry 287, 38936–38945. 10.1074/jbc.M112.415463.

45. Chen, T., Leng, J., Tan, J., Zhao, Y., Xie, S., Zhao, S., Yan, X., Zhu, L., Luo, J., Kong, L., et al. (2023). Discovery of Novel Potent Covalent Glutathione Peroxidase 4 Inhibitors as Highly Selective Ferroptosis Inducers for the Treatment of Triple-Negative Breast Cancer. J. Med. Chem. 66, 10036–10059. 10.1021/acs.jmedchem.3c00967.

46. Shintoku, R., Takigawa, Y., Yamada, K., Kubota, C., Yoshimoto, Y., Takeuchi, T., Koshiishi, I., and Torii, S. (2017). Lipoxygenase-mediated generation of lipid peroxides enhances ferroptosis induced by erastin and RSL3. Cancer Science 108, 2187–2194. 10.1111/cas.13380.

47. Sui, X., Zhang, R., Liu, S., Duan, T., Zhai, L., Zhang, M., Han, X., Xiang, Y., Huang, X., Lin, H., et al. (2018). RSL3 Drives Ferroptosis Through GPX4 Inactivation and ROS Production in Colorectal Cancer. Front. Pharmacol. 9. 10.3389/fphar.2018.01371.

48. Skouta, R., Dixon, S.J., Wang, J., Dunn, D.E., Orman, M., Shimada, K., Rosenberg, P.A., Lo, D.C., Weinberg, J.M., Linkermann, A., et al. (2014). Ferrostatins Inhibit Oxidative Lipid Damage and Cell Death in Diverse Disease Models. J. Am. Chem. Soc. 136, 4551–4556. 10.1021/ja411006a.

49. Dixon, S.J., Lemberg, K.M., Lamprecht, M.R., Skouta, R., Zaitsev, E.M., Gleason, C.E., Patel, D.N., Bauer, A.J., Cantley, A.M., Yang, W.S., et al. (2012). Ferroptosis: An Iron-Dependent Form of Nonapoptotic Cell Death. Cell 149, 1060–1072. 10.1016/j.cell.2012.03.042.

50. Yang, W.S., SriRamaratnam, R., Welsch, M.E., Shimada, K., Skouta, R., Viswanathan, V.S., Cheah, J.H., Clemons, P.A., Shamji, A.F., Clish, C.B., et al. (2014). Regulation of Ferroptotic Cancer Cell Death by GPX4. Cell 156, 317–331. 10.1016/j.cell.2013.12.010.

51. Hattori, K., Ishikawa, H., Sakauchi, C., Takayanagi, S., Naguro, I., and Ichijo, H. (2017). Cold stress-induced ferroptosis involves the ASK1-p38 pathway. EMBO Rep 18, 2067– 2078. 10.15252/embr.201744228.

52. Doll, S., Freitas, F.P., Shah, R., Aldrovandi, M., Da Silva, M.C., Ingold, I., Goya Grocin, A., Xavier Da Silva, T.N., Panzilius, E., Scheel, C.H., et al. (2019). FSP1 is a glutathione-independent ferroptosis suppressor. Nature 575, 693–698. 10.1038/s41586-019-1707-0.

53. Bersuker, K., Hendricks, J.M., Li, Z., Magtanong, L., Ford, B., Tang, P.H., Roberts, M.A., Tong, B., Maimone, T.J., Zoncu, R., et al. (2019). The CoQ oxidoreductase FSP1 acts parallel to GPX4 to inhibit ferroptosis. Nature 575, 688–692. 10.1038/s41586-019-1705-2.

54. Hu, Q., Wei, W., Wu, D., Huang, F., Li, M., Li, W., Yin, J., Peng, Y., Lu, Y., Zhao, Q., et al. (2022). Blockade of GCH1/BH4 Axis Activates Ferritinophagy to Mitigate the Resistance of Colorectal Cancer to Erastin-Induced Ferroptosis. Front Cell Dev Biol 10, 810327. 10.3389/fcell.2022.810327.

55. Ma, T., Du, J., Zhang, Y., Wang, Y., Wang, B., and Zhang, T. (2022). GPX4-independent ferroptosis—a new strategy in disease’s therapy. Cell Death Discov. 8, 1–8. 10.1038/s41420-022-01212-0.

56. Zhang, W., Dai, J., Hou, G., Liu, H., Zheng, S., Wang, X., Lin, Q., Zhang, Y., Lu, M., Gong, Y., et al. (2023). SMURF2 predisposes cancer cell toward ferroptosis in GPX4-independent manners by promoting GSTP1 degradation. Molecular Cell 83, 4352–4369.e8. 10.1016/j.molcel.2023.10.042.

57. Sone, M., Mitsuhashi, N., Sugiura, Y., Matsuoka, Y., Maeda, R., Yamauchi, A., Okahashi, R., Yamashita, J., Sone, K., Enju, S., et al. (2023). Identification of genes supporting cold resistance of mammalian cells: lessons from a hibernator. Preprint at bioRxiv, 10.1101/2023.12.27.573489 https://doi.org/10.1101/2023.12.27.573489.

58. Nakamura, T., Ogawa, M., Kojima, K., Takayanagi, S., Ishihara, S., Hattori, K., Naguro, I., and Ichijo, H. (2021). The mitochondrial Ca2+ uptake regulator, MICU1, is involved in cold stress-induced ferroptosis. EMBO Rep 22, EMBR202051532. 10.15252/embr.202051532.

59. Davey, H.M., and Hexley, P. (2011). Red but not dead? Membranes of stressed Saccharomyces cerevisiae are permeable to propidium iodide. Environ Microbiol 13, 163–171. 10.1111/j.1462-2920.2010.02317.x.

60. Kirchhoff, C., and Cypionka, H. (2017). Propidium ion enters viable cells with high membrane potential during live-dead staining. Journal of Microbiological Methods 142, 79–82. 10.1016/j.mimet.2017.09.011.

61. Avelar-Freitas, B.A., Almeida, V.G., Pinto, M.C.X., Mourão, F.A.G., Massensini, A.R., Martins-Filho, O.A., Rocha-Vieira, E., and Brito-Melo, G.E.A. (2014). Trypan blue exclusion assay by flow cytometry. Braz J Med Biol Res 47, 307–315. 10.1590/1414-431X20143437.

62. Cooper, S.T., Sell, S.S., Fahrenkrog, M., Wilkinson, K., Howard, D.R., Bergen, H., Cruz, E., Cash, S.E., Andrews, M.T., and Hampton, M. (2016). Effects of hibernation on bone marrow transcriptome in thirteen-lined ground squirrels. Physiol Genomics 48, 513–525. 10.1152/physiolgenomics.00120.2015.

63. Vermillion, K.L., Anderson, K.J., Hampton, M., and Andrews, M.T. (2015). Gene expression changes controlling distinct adaptations in the heart and skeletal muscle of a hibernating mammal. Physiol Genomics 47, 58–74. 10.1152/physiolgenomics.00108.2014.

64. Hampton, M., Melvin, R.G., and Andrews, M.T. (2013). Transcriptomic analysis of brown adipose tissue across the physiological extremes of natural hibernation. PLoS One 8, e85157. 10.1371/journal.pone.0085157.

65. Schwartz, C., Hampton, M., and Andrews, M.T. (2013). Seasonal and regional differences in gene expression in the brain of a hibernating mammal. PLoS One 8, e58427. 10.1371/journal.pone.0058427.

66. Yang, Y., Hao, Z., An, N., Han, Y., Miao, W., Storey, K.B., Lefai, E., Liu, X., Wang, J., Liu, S., et al. (2023). Integrated transcriptomics and metabolomics reveal protective effects on heart of hibernating Daurian ground squirrels. J Cell Physiol 238, 2724–2748. 10.1002/jcp.31123.

67. Li, F.-J., Long, H.-Z., Zhou, Z.-W., Luo, H.-Y., Xu, S.-G., and Gao, L.-C. (2022). System Xc -/GSH/GPX4 axis: An important antioxidant system for the ferroptosis in drug-resistant solid tumor therapy. Front Pharmacol 13, 910292. 10.3389/fphar.2022.910292.

68. Forman, H.J., Zhang, H., and Rinna, A. (2009). Glutathione: Overview of its protective roles, measurement, and biosynthesis. Mol Aspects Med 30, 1–12. 10.1016/j.mam.2008.08.006.

69. Zheng, J., and Conrad, M. (2020). The Metabolic Underpinnings of Ferroptosis. Cell Metabolism 32, 920–937. 10.1016/j.cmet.2020.10.011.

70. Doll, S., Proneth, B., Tyurina, Y.Y., Panzilius, E., Kobayashi, S., Ingold, I., Irmler, M., Beckers, J., Aichler, M., Walch, A., et al. (2017). ACSL4 dictates ferroptosis sensitivity by shaping cellular lipid composition. Nat Chem Biol 13, 91–98. 10.1038/nchembio.2239.

71. Asghar, W., El Assal, R., Shafiee, H., Anchan, R.M., and Demirci, U. (2014). Preserving human cells for regenerative, reproductive, and transfusion medicine. Biotechnology Journal 9, 895–903. 10.1002/biot.201300074.

72. Yoo, S.-E., Chen, L., Na, R., Liu, Y., Rios, C., Remmen, H.V., Richardson, A., and Ran, Q. (2012). Gpx4 ablation in adult mice results in a lethal phenotype accompanied by neuronal loss in brain. Free Radic Biol Med 52, 1820–1827. 10.1016/j.freeradbiomed.2012.02.043.

73. Concordet, J.-P., and Haeussler, M. (2018). CRISPOR: intuitive guide selection for CRISPR/Cas9 genome editing experiments and screens. Nucleic Acids Res 46, W242– W245. 10.1093/nar/gky354.

74. Stewart, S.A., Dykxhoorn, D.M., Palliser, D., Mizuno, H., Yu, E.Y., An, D.S., Sabatini, D.M., Chen, I.S.Y., Hahn, W.C., Sharp, P.A., et al. (2003). Lentivirus-delivered stable gene silencing by RNAi in primary cells. RNA 9, 493–501. 10.1261/rna.2192803.

75. Langmead, B., Trapnell, C., Pop, M., and Salzberg, S.L. (2009). Ultrafast and memory-efficient alignment of short DNA sequences to the human genome. Genome Biol 10, R25. 10.1186/gb-2009-10-3-r25.

76. Li, W., Xu, H., Xiao, T., Cong, L., Love, M.I., Zhang, F., Irizarry, R.A., Liu, J.S., Brown, M., and Liu, X.S. (2014). MAGeCK enables robust identification of essential genes from genome-scale CRISPR/Cas9 knockout screens. Genome Biol 15, 554. 10.1186/s13059-014-0554-4.

77. R Core Team (2022). R: A Language and Environment for Statistical Computing | BibSonomy. R Foundation for Statistical Computing.

78. Wei T, S.V. (2021). R package “corrplot”: Visualization of a Correlation Matrix (Version 0.92). https://github.com/taiyun/corrplot.

79. Silva, D., Santos, G., Barroca, M., and Collins, T. (2017). Inverse PCR for Point Mutation Introduction. In PCR: Methods and Protocols Methods in Molecular Biology., L. Domingues, ed. (Springer), pp. 87–100. 10.1007/978-1-4939-7060-5_5.

80. Schulte, F., Hasturk, H., and Hardt, M. (2019). Mapping Relative Differences in Human Salivary Gland Secretions by Dried Saliva Spot Sampling and nanoLC–MS/MS. Proteomics 19, 1900023. 10.1002/pmic.201900023.

